# Molecular and functional diversity of distinct subpopulations of extracellular vesicles from stressed pancreatic beta cells: implications for autoimmunity

**DOI:** 10.1101/2020.03.26.003145

**Authors:** Khem Raj Giri, Laurence de Beaurepaire, Dominique Jegou, Margot Lavy, Mathilde Mosser, Aurelien Dupont, Romain Fleurisson, Laurence Dubreil, Mayeul Collot, Peter Van Endert, Jean-Marie Bach, Gregoire Mignot, Steffi Bosch

## Abstract

Beta cell failure and apoptosis following islet inflammation have been associated with autoimmune type 1 diabetes pathogenesis. As conveyors of biological active material, extracellular vesicles (EV) act as mediators in communication with immune effectors fostering the idea that EV from inflamed beta cells may contribute to autoimmunity. Evidence accumulates that beta exosomes promote diabetogenic responses, but relative contributions of larger vesicles as well as variations in the composition of the beta cell’s vesiculome due to environmental changes have not been explored yet. Here, we made side-by-side comparisons of the phenotype and function of apoptotic bodies (AB), microvesicles (MV) and small EV (sEV) isolated from an equal amount of MIN6 beta cells exposed to inflammatory, hypoxic or genotoxic stressors. Under normal conditions, large vesicles represent 93% of the volume, but only 2% of the number of the vesicles. Our data reveal a consistently higher release of AB and sEV and to a lesser extent of MV, exclusively under inflammatory conditions commensurate with a 4-fold increase in the total volume of the vesiculome and enhanced export of immune-stimulatory material including the autoantigen insulin, microRNA, and cytokines. Whilst inflammation does not change the concentration of insulin inside the EV, specific Toll-like receptor-binding microRNA sequences preferentially partition into sEV. Exposure to inflammatory stress engenders drastic increases in the expression of monocyte chemoattractant protein 1 in all EV and of interleukin-27 solely in AB suggesting selective sorting towards EV subspecies. Functional *in vitro* assays in mouse dendritic cells and macrophages reveal further differences in the aptitude of EV to modulate expression of cytokines and maturation markers. These findings highlight the different quantitative and qualitative imprints of environmental changes in subpopulations of beta EV that may contribute to the spread of inflammation and sustained immune cell recruitment at the inception of the (auto-) immune response.

**Graphical Abstract:** 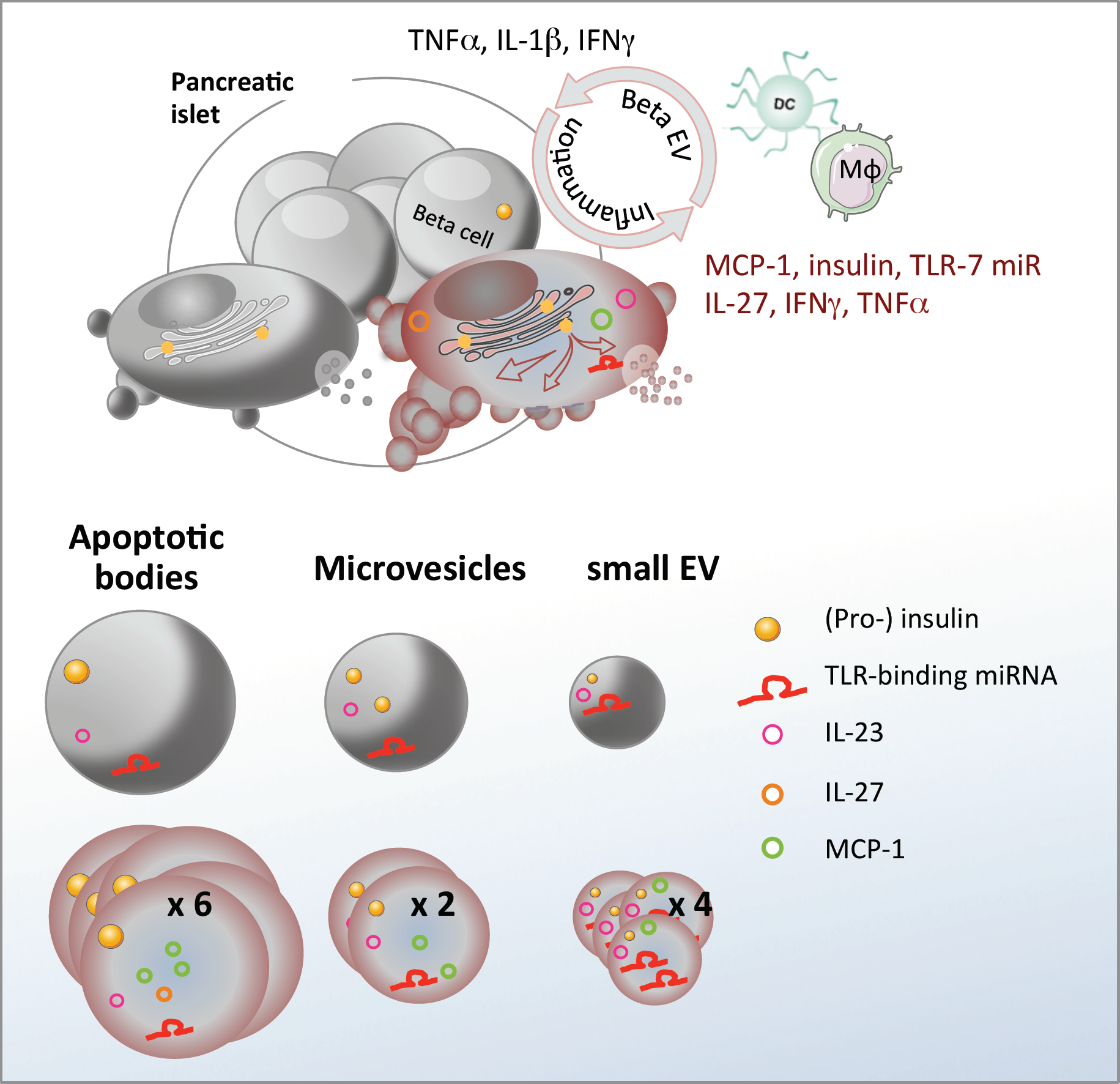

Inflammation stimulates release of a heterogeneous population of beta EV with differential expression of immunogenic substances involved in immune cell recruitment and activation.

**Highlights:**

- Stress engenders an up to four-fold increase in the volume of the vesiculome and enhanced auto-antigen release
- Cytokines are selectively sorted into EV subspecies
- TLR-binding microRNAs are enriched in sEV
- EV from stressed beta cells promote dendritic and macrophage cell activation

## Introduction

Type 1 diabetes (T1D) is an autoimmune disease caused by the destruction of the insulin-producing beta cells in the pancreas leading to chronic hyperglycaemia and serious long-term complications such as cardiovascular disease, neuropathy, nephropathy and blindness (reviewed in ^1^). More than 30 million of people suffer from T1D worldwide (www.idf.org). T1D and its sequelae reduce life expectancy of patients by more than eleven years ^2^. Pathogenesis of T1D is characterized by inflammatory events in the beta cell microenvironment causing innate immune activation followed by progressive infiltration of the islets of Langerhans in the endocrine pancreas by auto-reactive cytotoxic T-lymphocytes. Disease aetiology has only partially been elucidated, but results from a complex interplay between genetic and environmental factors collectively engendering functional defects in the immune system and the beta cell itself. Environmental changes in toxins, pathogens, nutrients in particular glucose overload, and low physical activity have been suggested to be responsible for the 3.4% annual increase in disease incidence ^3^. Due to its demanding secretory function, the beta cell is extremely sensitive to stress. Insulin accounts for up to half of the cell’s protein content ^4^ and rapid changes can exceed the endoplasmic reticulum’s (ER) folding capacities leading to the accumulation of misfolded proteins e.g. potential neoantigens within the lumen of the ER. By interaction with built-in sensors, these misfolded proteins trigger the unfolded protein response (UPR), a signalling pathway that aims to restore homeostasis by enhancing the cell’s folding capacity and translational attenuation. However, chronic stress can cause the UPR to initiate apoptosis. Beta cell stress and apoptosis has been associated with T1D pathogenesis ^5,6^, yet, how stressed beta cells trigger innate immune responses at disease initiation has not been fully elucidated.

Extracellular vesicles (EV) are membrane-bound vesicles released by healthy and diseased cells. Three major types of EV can be distinguished based on their origin and biogenesis pathway: apoptotic bodies (AB), microvesicles (MV) and exosomes (reviewed in ^7^). AB are large 1,000 - 5,000 nm vesicles released by cells undergoing apoptosis ^8^. In contrast to other EV, AB may contain cellular organelles and their constituents including elements from the nucleus, mitochondria, the Golgi apparatus, the ER and the cytoskeleton. MV are formed by outward budding and scission of the plasma membrane. Typically, the size of MV ranges from 100-1,000 nm. In line with their pathway of formation, MV contain mainly cytosolic and plasma membrane-associated proteins such as tetraspanins. Exosomes result from the inward budding of the membrane of the endosome leading to the formation of 30-120 nm intraluminal vesicles that can be released upon fusion of the endosome with the plasma membrane. Throughout its maturation, the endosome is an important site of bi-directional translocation of substances between the cytoplasm and the endosome. In consequence, the packing of specific cargo molecules into EV and their release are intimately linked to the state of the releasing cells. As conveyors of biological active material from their cell of origin to neighbouring or distant recipient cells, EV act as mediators in cell-to-cell communication fostering the idea that they may constitute the missing link between beta cell stress and immune activation (reviewed in ^9^).

While beta AB have been successfully used to induce tolerance in diabetes-prone non obese diabetic (NOD) mice ^10^, evidence accumulates that exosomes derived from pancreatic beta cells contribute to T1D development. Strikingly, all known beta auto-antigens are directly or indirectly linked to secretory pathways and thirteen localize to secretory granules, synaptic vesicles, the ER and the trans-Golgi network (TGN)^11^. Several studies showed that human and mouse beta EV contain major auto-antigens of type 1 diabetes such as glutamic acid decarboxylase 65 (GAD65), glucose-transporter 2 (GLUT-2), islet-associated antigen 2 (IA-2), zinc transporter 8 (Znt8), and insulin ^12-15^. Interconnections between the TGN secretory pathway of autoantigens and the endosomal compartment where exosome biogenesis occurs have convincingly been demonstrated by immunofluorescent studies co-localising the auto-antigen GAD65 with the TGN protein 38, but also with endosomal ras-related protein in brain 11 (Rab11) and the exosomal markers flotillin-1 (FLOT1) and CD81 in vesicular structures at the peripheral membrane ^12^. Exosomes from healthy beta cells efficiently trigger antigen-presenting cell (APC) activation and T-cell proliferation *in vitro* and accelerate islet infiltration by immune cells in non obese diabetic resistant mice *in vivo*^13^. In human T1D patients, healthy beta EV mediate B- and T-cell activation^16^. It has further been hypothesized that aberrant sorting in stressed beta cells could fuel release of misfolded immunogenic proteins and danger-associated molecular patterns (DAMP) inside EV. With the aim to explore roles of beta EV in T1D pathogenesis, attempts are made to recreate the beta cell environment by adding a mild cocktail of the proinflammatory cytokines TNFα, IFNγ and IL-1β present in the pancreas at disease initiation^12,17,18^. EV ferry short non-coding microRNA (miRNA) that have the aptitude to repress translation of target genes in recipient cells^19^, a well-documented mechanism termed RNA interference (RNAi)^20^. MiRNA in exosomes derived from beta cells under inflammatory conditions contribute to the spread of beta cell apoptosis ^17^. However, the biological relevance of miRNA transfer has been questioned by estimates of 1,000 copies required per recipient cells to allow for effective target gene regulation^21^. More recently, six specific GU-rich miRNA sequences have been identified (let-7b/c, miR-21, miR-7a, miR-29a/b) that may stimulate immune signalling by binding to the Toll-like receptor-7 (TLR-7), independently of RNAi. Packed into EV, these miRNA sequences act as DAMP exacerbating inflammation in cancer, neurological and autoimmune settings ^22-28^. Beta EV T- and B-cell activation in NOD mice was impaired in NOD.*MyD88*^*-/-*^ mice suggesting a role for TLR-signalling in EV-mediated immune responses^13,29^.

To date, the molecular and functional diversity of EV in the beta cell’s secretome has not been thoroughly explored. The majority of beta EV studies focuses on small exosome-like vesicles and AB and no studies on contributions of beta MV have been published to our knowledge. Because subtypes of beta EV potentially exert detrimental or protective effects in the immune balance, side-by-side comparisons are mandatory to evaluate their role in T1D pathogenesis. We herein sought to investigate on changes in the relative composition of the vesiculome as well as the partition of the candidate autoantigen insulin and immunostimulatory miRNA sequences inside AB, MV and exosome subpopulations derived from equal amounts of healthy and stressed beta cells and their impact on innate immune responses. As current isolation methods do not allow distinguishing between exosomes of endosomal origin and small MV, the later will be called small EV (sEV) throughout this study.

## Materials and methods

### Mice

NOD/ShiLtJ mice were obtained from Charles River Laboratories (L’Abresle, France), bred and housed in a pathogen-free environment at ONIRIS’ Rodent Facility (Agreement #44266). Six to ten weeks old female mice were used in the study. All animal procedures were approved by the Pays de la Loire regional committee on ethics of animal experiments (APAFIS#9871). All possible efforts were made to minimize animal suffering.

### Cell culture

MIN6 cells (kindly provided by Prof. Jun-ichi Miyazaki, University Medical School, Osaka, Japan) were cultured at a density of 1.5 × 10^5^ cells/cm^2^ in DMEM high glucose medium (Life Technologies, Saint Aubin, France) supplemented with 10% FCS (Eurobio, Les Ulis, France) and 20 µM beta-mercaptoethanol (SIGMA, Saint Quentin Fallavier, France)^30^. Cell cultures were regularly assessed for mycoplasma contamination using the mycoplasma quick test (Lonza, Basel, Switzerland). For EV production, MIN6 cells were plated at a density of 3 × 10^5^ cells/cm^2^ in DMEM medium. The following day, the medium was replaced by OptiMEM (Life Technologies) supplemented with 1% exosome-precleared FCS (FCS^exo-^) obtained through overnight centrifugation at 120,000 × g on a SW41 Ti swinging bucket rotor on a L7-55 centrifuge (Beckman Coulter, Villepinte, France) in polyallomer tubes (Beckman Coulter). For experimental induction of cellular stress, MIN6 cells were either exposed to pro-inflammatory cytokines (17U/mL IL-1β, 167U/mL TNFα and 17U/mL IFNγ; all cytokines were supplied by eBioscience Affymetrix, Paris, France) 254 nm UVB irradiation 10 mJ, BLX-254, Vilber Lourmat, Marne la Vallee, France), low oxygen tension (1% O_2_) or left untreated. Supernatants containing MIN6 exosomes were harvested 30 hours later. At harvest, cell viability of untreated cells was ≥ 90% in compliance with guidelines to minimize apoptotic EV input in vesicle preparations ^31,32^.

RAW264.7 cells (ATCC #TIB-71) were maintained in RPMI 1640 medium (Life Technologies), supplemented with 10% FCS and 2 mM L-glutamine (Eurobio). Cells were split twice per week by gently scrapping of the cells in cold PBS 2mM EDTA solution. For immune assays, 5 × 10^4^ RAW264.7 cells were plated on flat-bottomed 96-well plates in RPMI 10% FCS^exo-^ medium 2 mM glutamine.

Bone marrow-derived dendritic cells (bmDC). Bone marrow progenitor cells were isolated from femurs and tibias from NOD_Shi/LTJ_ female mice and cultured in complete RPMI medium (Eurobio) i.e. supplemented with 1% heat-denatured syngeneic mouse serum along with 1 mM sodium pyruvate, 100 IU/mL penicillin, 100 µg/mL streptomycin, 2 mM L-glutamine, non-essential amino acids and 20 µM beta-mercaptoethanol. Medium was supplemented with 20 ng/mL GM-CSF (PeproTech, Neuilly-sur-Seine, France) and 5 ng/mL IL4 (BioLegend, London, UK) and 2 × 10^6^ cells were cultured in 10mL of medium per 100 mm Petri dish for ten days. On day four and nine, an additional 10 mL and 5 mL of complete culture media were added, respectively. On day 7, 10 mL of culture media were refreshed. On day ten, flow cytometry routinely revealed CD11c^+^ > 90% purity of bmDC cultures.

For immune assays, 0.3 × 10^6^ bmDC were cultured in 48-well plates coated polyhydroxyethylmethacrylate (Sigma) for eighteen hours in RPMI 1% exosome-depleted mouse serum (MS^exo-^) supplemented with additives as described before. EV or Toll-like receptor–ligands (TLR-L) (InvivoGen, Toulouse, France): imiquimod (IMIQ; TLR-7L), resiquimod (R848; TLR-7/8L) and polyinosine-polycytidylic acid (PIC; TLR-3L) were used at concentrations as indicated on figures.

### Antibodies & reagents

Phenotypic analysis was performed by flow cytometry on a FACS Aria (BD Biosciences, Le Pont de Claix, France) or MACSQuant (Miltenyi, Paris, France) instrument using the following antibodies: CD11c (N418, BioLegend, Toulouse, France), CD86 (GL-1, BioLegend), major histocompatibility complex II (MHC II; 103.6.2, BD Biosciences), CD40 (3/23, BioLegend), CD115 (CSF-1R, BioLegend), CD11b (M1/70, BioLegend), B220 (RA3-6B2, BD Biosciences). Zombie-NIR (BioLegend), DAPI (Fisher Scientific, Illkirch, France) or Viobility 405/520 (Miltenyi) dyes were used to discriminate death cells. Results were analysed using FlowJo (Tree Star Inc., Ashland, OR, USA) or Flowlogic (Miltenyi) software.

Cytokine secretion into cell culture supernatants was quantified by ELISA using IFNa, TNFα (R&D systems) according to the suppliers’ protocols. Cytometric bead assay analysis of IL-23, IL-1α, IFNγ, TNFα, MCP-1, IL-12p70, IL-1β, IL-10, IL-6, IL-27, IL-17A, IFNβ and GM-CSF was performed using a predefined 13-plex Mouse Inflammation panel (BioLegend). To assess for internal as well as surface-associated cytokines, 30 minutes of incubation in 0.5% Triton X-100 and 1 min. of sonication were performed prior to CBA of cytokine expression in EV. Total mouse insulin in MIN6 EV was quantified by ELISA (Mercodia, Upsala, Sweden).

Protein concentration was determined by a Bradford protein assay using Coomassie plus assay reagent (Fisher Scientific). Optical densities were read on Fluostar (BMG LABTECH, Champigny sur Marne, France) and Nanodrop2000 (Fisher Scientific) spectrophotometers following the supplier’s recommendations.

### Markers of hypoxia

For detection of the endogenous marker of hypoxia HIF-1α, MIN6 cells were cultured as indicated for 30h in normoxia or hypoxia. As soon as the hypoxia chamber was opened, cells were washed and lysed with 1mL of 50mM Tris (pH 7.4), 300 mM NaCl, 10% (w/v) glycerol, 3 mM EDTA, 1 mM MgCl2, 20 mM b-glycerophosphate, 25 mM NaF, 1% Triton X-100, 25 µg/mL Leupeptin, 25 µg/mL Pepstatin and 3µg/mL Aprotinin lysis buffer for 30 min on a rotating platform at 4°C. The cells were centrifuged at 2,000 × g for 5 min and supernatants were stored at -80°C. Proteins were quantified using the Bradford assay and HIF-1α was measured by ELISA following the supplier’s instructions (R&Dsystem). Optical Density at 450nm was measured using the FLUOstar Optima Microplate Reader (BMG Labtech, Champigny sur Marne, France).

The exogenous marker pimonidazole forms stable adducts with thiol groups in cellular proteins under hypoxia. Briefly, 3×10^5^ cells/cm^2^ MIN6 were cultured for 24 hours in DMEM culture medium. After washing, cells were incubated with 100 µM pimonidazole (Hypoxyprobe-1 plus; Hypoxyprobe Inc, Burlington, USA) in OptiMEM 1% FCS^exo-^ for 2 hours in normoxia or hypoxia. The cells were fixed with 4% PFA, permeabilized with PBS 0.1% Triton 4% FCS followed by saturation in PBS 5% rabbit serum for 1 hour. For detection of pimonidazole adducts, cells were incubated overnight at 4°C with FITC-conjugated mouse anti-pimonidazole monoclonal antibody (1:100). The cells were washed and imaged on a fluorescent microscope (AXIO Zeiss, Leica, Nanterre, France).

### Caspase assay

Apoptosis was assayed by fluorescent caspase-3/7 substrate cleavage staining. Briefly, 3×10^5^ cells/cm^2^ MIN6 cells were cultured in eight-chamber labteks with coverslips. After overnight culture, cells were switched to OptiMEM 1% FCS production medium and exposed to cytokines, UV irradiation, hypoxia or left untreated. Eighteen hours later, cells were treated with 2 mM caspase-3/7 detection reagent (Fisher Scientific) for 30 min. at 37°C and counterstained with 1 µg/ml Hoechst 33342 (Sigma). Cells were fixed with 4% PFA, washed with PBS and overlaid with Mowiol (Sigma) before analysis by fluorescence confocal imaging on a LSM780 confocal microscope (Zeiss, Oberkochen, Germany). Tiles of nine images per well were acquired and processed for semi-automatic quantitative analysis of caspase-positive cells using an in-house macro and Fiji software. Total cell count was set equal to the number of Hoechst positive regions.

### Separation of beta-EV subpopulations

EV were collected from MIN6 supernatants using a method combining differential centrifugation, ultrafiltration and size-exclusion chromatography steps. Briefly, 90 ml of 30-hours supernatants from MIN6 cells were centrifuged immediately after harvest at 300 × *g* 10 min., 2,000 × g, 20 minutes (AB) and 16,600 × g, 20 minutes (MV). The pellets containing AB and MV were washed with PBS or RPMI and centrifuged again before use. The 16,500 supernatants were filtered 0.2µm and concentrated on an AMICON MWCO-100 kDa cellulose ultrafiltration unit (Dutscher, Issy-les-Moulineaux, France). Approximately, 100 µl of concentrates were recovered and passed through a size exclusion chromatography column (IZON, Lyon, France). sEV were collected following the supplier’s recommendations in flow-through fractions four and eight for qEV single and qEV original, respectively. EV were stored at 4 °C for one to three days or at -80 °C for up to one year. All assays of biological activity were carried out using fresh EV. For transcriptomic analyses, 90 ml of 30-hours supernatants from MIN6 cells were centrifuged immediately after harvest at 300 × *g* 10 min., followed by centrifugation at 16,500 × g, 20 minutes to collect large EV (LEV) comprising both AB and MV.

### Tunable resistive pulse sensing (TRPS)

The size and concentration of EV were analysed by the TRPS technique using a qNANO instrument and NP2000 (AB), NP800 (MV) and NP100 (sEV) nanopores (IZON, Lyon, France). All samples were diluted in PBS 0.03% Tween-20. After instrument calibration using 110 nm, 710 nm or 2000 nm calibration beads (Izon), all samples were recorded with at least two different pressures. Respective particle volumes V=4/3πr^3^ x nb particles were calculated based on the mean particle diameter measured, assuming spherical shape.

### Immunoblotting

Protein lysats of cells, AB and MV were prepared in RIPA buffer containing a cocktail of protease inhibitors (Sigma). Proteins were denatured in Laemmli buffer and separated by 4-12% gradient SDS-PAGE in non-reducing (tetraspanins CD81, CD63, CD9) or reducing (all other) conditions and transferred to a nitrocellulose membrane (CD63, flotillin-1, peIF2a and CHOP; Fisher Scientific) or PVDF membrane (all other; BIO-RAD, Marnes La Coquette, France). Membranes were incubated with primary antibodies CD63 (NVG-2; 1:1000; BioLegend), CD81 (Eat-2; 1:1000; BioLegend), CD9 (EM-04; 1:1000; Abcam, Cambridge, UK), polyclonal rabbit anti-calnexin antibody (1:1000; Euromedex, Souffelweyersheim, France), β–actin (W16197A; 1:20,000; Biolegend), flotillin-1 (W16108A; 1:1000; Biolegend), peIF2a (D9G8; 1:1000; Ozyme) and CHOP (D46F1; 1:1000; Ozyme) blocked by either TBS 0.05% Tween-20 4% BSA (CD81, peIF2a, CHOP) or TBS 0.05% Tween-20 5% milk (CD9, CD63, calnexin, flotillin-1), followed by incubation with cognate HRP-conjugated secondary antibodies 1:100,000. The signals were detected with enhanced chemiluminescence substrate (ECL West Pico Femto, Fischer Scientific) on a Fusion FX6 instrument (Fisher Scientific).

### Cryo-electron microscopy

MV and sEV were applied onto glow-discharged perforated grids (C-flat™), prepared using an EM-GP (Leica, Germany) at room temperature in a humidity saturated atmosphere. EV samples were mixed with 10 nm diameter gold nanoparticles at a concentration of 80nM^33^ and four µl of the mixture were deposited on the grids. Excess sample was removed by blotting for 0.8 to 1.2 seconds before snap-freezing of samples into liquid ethane and storage in liquid nitrogen until observation. The grids were mounted in a single-axis cryo-holder (model 626, Gatan, USA) and the data were collected on a Tecnai G^2^T20 sphera electron microscope (FEI company, The Netherlands) equipped with a CCD camera (US4000, Gatan) at 200 kV. Images were taken at a nominal magnification x 29,000 in low-electron dose conditions. For cryo-electron tomography, single-axis tilt series, typically in the angular range ±60°, were acquired under low electron doses (∼0.3 e^-^/Å^2^) using the camera in binning mode 2 and at nominal magnifications of 25,000x and 29,000x, corresponding to calibrated pixel sizes of 0.95 and 0.79 nm at the specimen level, respectively. Tomograms were reconstructed using the graphical user interface eTomo from the IMOD software package ^34,35^.

### Fluorescence imaging of EV

AB and MV were separated from MIN6 cell supernatant and stained with MemBright (MB) dye (MB-Cy3 and MB-Cy5 (200 nM) kindly provided by M Collot). The mixture was incubated for 30 minutes at room temperature with gentle rotation. EV were then centrifuged at 2,000 × g (AB) or 16,600 × g (MV) for 20 minutes and washed in PBS. Samples were transferred into Labtek wells and overlaid with Mowiol (Sigma) before acquisition of images in superresolution mode Airyscan on a Zeiss LSM780 instrument.

### Quantitative RT-PCR analysis

Total RNA including miRNA was extracted from MIN6 cells or EV derived from an equal amount of cells using the miRVana kit (Fisher Scientific) or TriReagent (SIGMA), respectively. During the initial lysis step, all samples were spiked with 10^10^ copies of a synthetic analogue of ath-miR-159 (Eurogentec, Angers, France). Following reverse transcription using RT-stem-loop primers, extravesicular cDNA was pre-amplified for all miRNA except the spike ath-miR-159 by 10-14 cycles of PCR using Taqman probe reagent (Solisbiodyne, Tartu, Estonia) and Taqman assays (Fisher Scientific), followed by 40 cycles of PCR on an ABI7300 instrument (Fisher Scientific). For each target, standard curves were generated using serial sample dilutions. Relative quantities (in arbitrary units) of miRNA in samples were inferred by the relative standard curve method and normalised with respect to the spike and untreated controls.

### Statistical analysis

Statistical tests were performed using either Prism GraphPad Software (Comparex, Issy-les-Moulineaux, France) or R 3.6.0 ^36^ with RStudio ^37^ and lsmeans^38^ and lme4^39^ packages using tests as indicated in the figure legends. Confidence levels of 95% were considered significant. For linear mixed model, the parameters were the “type of EV” or the “treatment of the producing cells”. No interaction test was deemed necessary, as analysis was performed for all “treatments” of a single “type of EV” or vice-versa. The random parameter was the individual “experiment”. Post-hoc analysis was performed by the Tukey’s range test for pairwise comparisons on calculated least-square means.

## Results

Primary islet inflammatory events have been associated with beta cell stress and failure at the origin of T1D pathogenesis ^40,41^. With the aim to study the impact of cellular stress on the beta vesicular secretome, murine MIN6 beta cells were either left untreated (CTL) or exposed to a cocktail of mild doses of pro-inflammatory cytokines (CK) encountered at disease initiation. To discriminate between inflammation-specific and general responses to cellular stress, hypoxic (1% O_2_, HX) and genotoxic (ultraviolet irradiation, UV) stress situations were introduced (Fig.1A). Cells grown under hypoxia were assessed for the expression of endogenous and exogenous markers of hypoxia. Added to culture, pimonidazole hydrochloride forms adducts with thiol groups in proteins in cells at low oxygen tension (pO_2_ < 10 mmHg). Immunofluorescent microscopy analysis revealed the presence of pimonidazole adducts in hypoxic cells (Fig.1B). The hypoxia-inducible factor 1 (HIF-1) is a transcriptional regulator of the cellular response to low oxygen levels. Under hypoxic conditions, the subunit HIF-1α associates with the subunit HIF-1β and binds to the hypoxia response element (HRE) of target genes, initiating their expression ^42,43^. ELISA analysis showed a three-fold increase in HIF-1α expression from 9 (7-12) pg/mL (median (range)) for cells grown under normoxic conditions compared to 29 (15-44) pg/mL for cells grown under hypoxic conditions (Fig.1C; p = 0.0143).

**Fig1.**
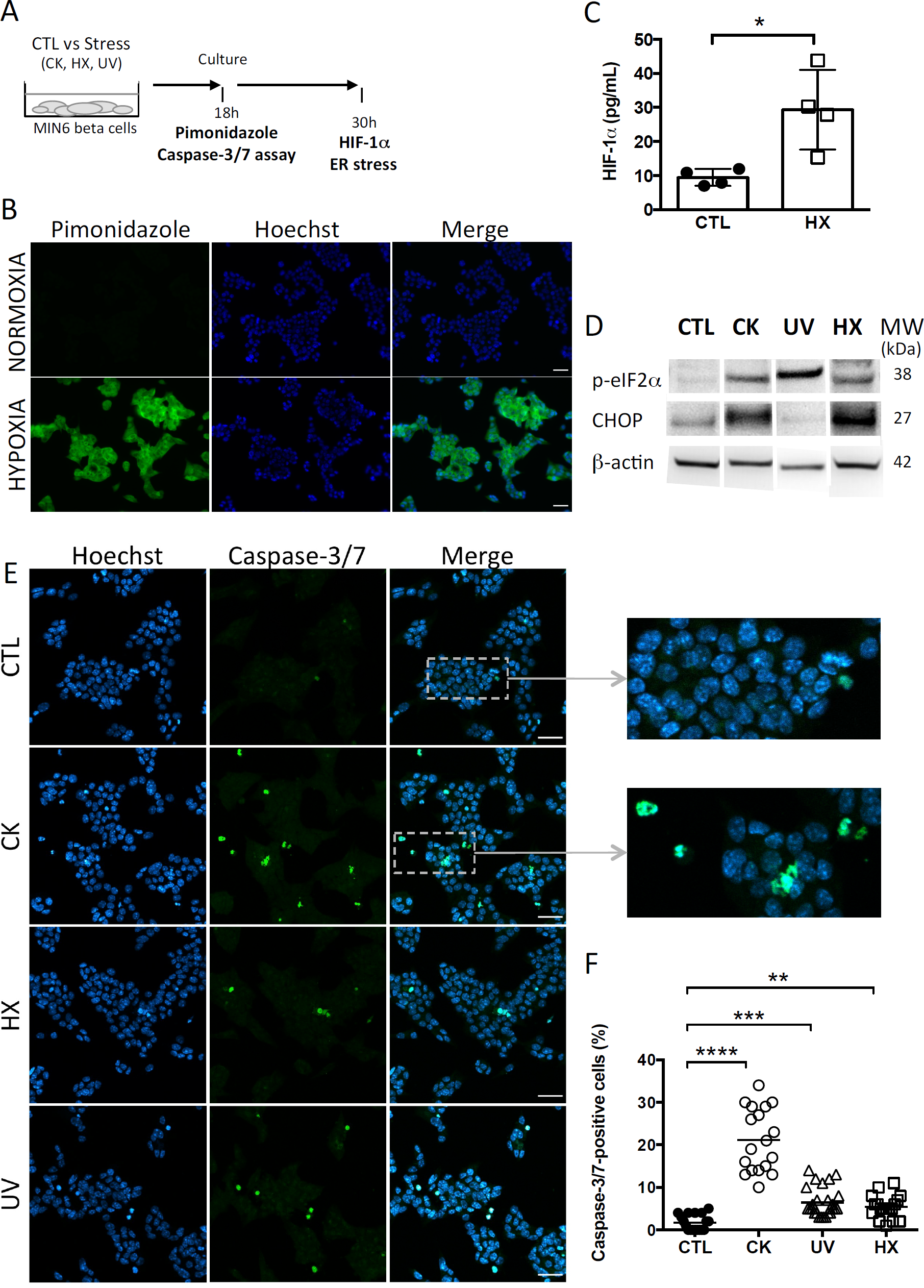
Experimental beta cell stress induces apoptosis. (A) After exposure to stress (TNFα, IL-1β, IFN-γ cytokines; CK), hypoxia (HX), ultraviolet (UV)) or not (control (CTL), MIN6 cells were cultured in 1% FCS exosome-depleted OptiMEM medium. (B-C) For cultures grown under normoxic and hypoxic conditions, (B) After 24h of culture, MIN6 cells were treated with 100µM pimonidazole and incubated for 2h followed by immunohistochemical detection of pimonidazole adducts (green). Nuclei were counterstained with Hoechst 33342 (blue). Scalebar 30 µm. Representative images of one out of three independent experiments are shown. (C) ELISA of the expression of HIF-1α in cells grown under normoxia or hypoxia for 30h. Data from 4 independent experiments are depicted with median and range. Mann-Whitney test, one-tailed, *p<0.05. (D) After 30h of culture, expression of markers of ER stress p-eIF2α and CHOP was analysed by western blotting and then the membranes were reprobed to β-actin. (E-F) After 18h of culture, the cells were stained with CellEvent Caspase-3/7 reagent (green) or Hoechst 33342 (nucleic dye, blue). (E) Fluorescence microscopy images of MIN6 cells reveal cells undergoing apoptosis as shown by the presence of caspase-3/7- positive cells. Scale bar 30µm. (F) The quantitative analysis showed a significant increase in the percentage of caspase-positive cells for treated in comparison to untreated control cells. Each data point represents results obtained for one image. Data are compiled from n=18 images (> 8000 nuclei) per situation from two independent experiments. Kruskal-Wallis test (**p<0.01, ***p<0.001 and ****p<0.0001).

Exposure to experimental stress induced ER stress as revealed by enhanced expression of the phosphorylated form of the subunit alpha of the eukaryotic translation initiation factor 2 (p-eIF-2α) and the transcription factor C/EBP homologous protein (CHOP), two effectors of the unfolded protein response (UPR) to ER stress (Fig.1D). While p-eIF-2α participates to translational attenuation with the aim to restore protein homeostasis in the ER at an early stage of the UPR, CHOP is activated belatedly after prolonged stress and controls cell fate by regulating expression of genes involved in apoptosis. After 30 hours of culture, cytokine- and HX-treated cells expressed CHOP in contrast to UV-irradiated cells. This difference might be explained by altered and presumably delayed kinetics of activation of the UPR following DNA damage at random by UV-light that pass through transcriptional and translational steps prior to changes in the proteome.

All treatment conditions engendered apoptosis in MIN6 cells as shown by the significant increases in the percentage of effector caspase-3/7- positive cells (Fig. 1E-F). The quantitative analysis of fluorescent caspase-substrate cleavage on confocal microscopy images (total > 8,000 nuclei counted for each situation) revealed a low percentage of 1.5 (0-5)% (median and (range)) of caspase-3/7-positive cells in untreated controls. Following exposure to stress, this percentage shifted to 20 (10-34)% for CK-, 5 (3-14)% for UV-, and 5 (1-11)% for HX-treated cells. Live cell imaging was performed to monitor the kinetics of apoptosis in individual cells (Suppl.Fig.1). In response to cytokines, caspase-3/7 activity appeared after five hours of treatment and steadily increased. Close to all cytokine-treated cells became apoptotic by the end of the 30h incubation period in contrast to untreated controls. Collectively, these results demonstrate that stress in our experimental conditions rapidly induces critical executioners of the cellular stress response in MIN6 beta cells cumulating in apoptosis.

To assess downstream effects of cellular stress on the beta cell’s secretome, AB, MV and sEV subpopulations were enriched from 30h conditioned MIN6 culture supernatants following a protocol combining differential centrifugation ^44^, (ultra-) filtration and size-exclusion chromatography ^45^ steps (outlined in Fig.2A). Western blot revealed the presence of vesicular markers i.e. the membrane proteins CD81, CD63, CD9 and flotillin, and the cytosolic protein beta-actin in all EV subpopulations (Fig.2B). The ER protein calnexin was present in AB and MV but absent in sEV in line with enrichment in vesicles of endosomal origin in the later.

**Fig2.**
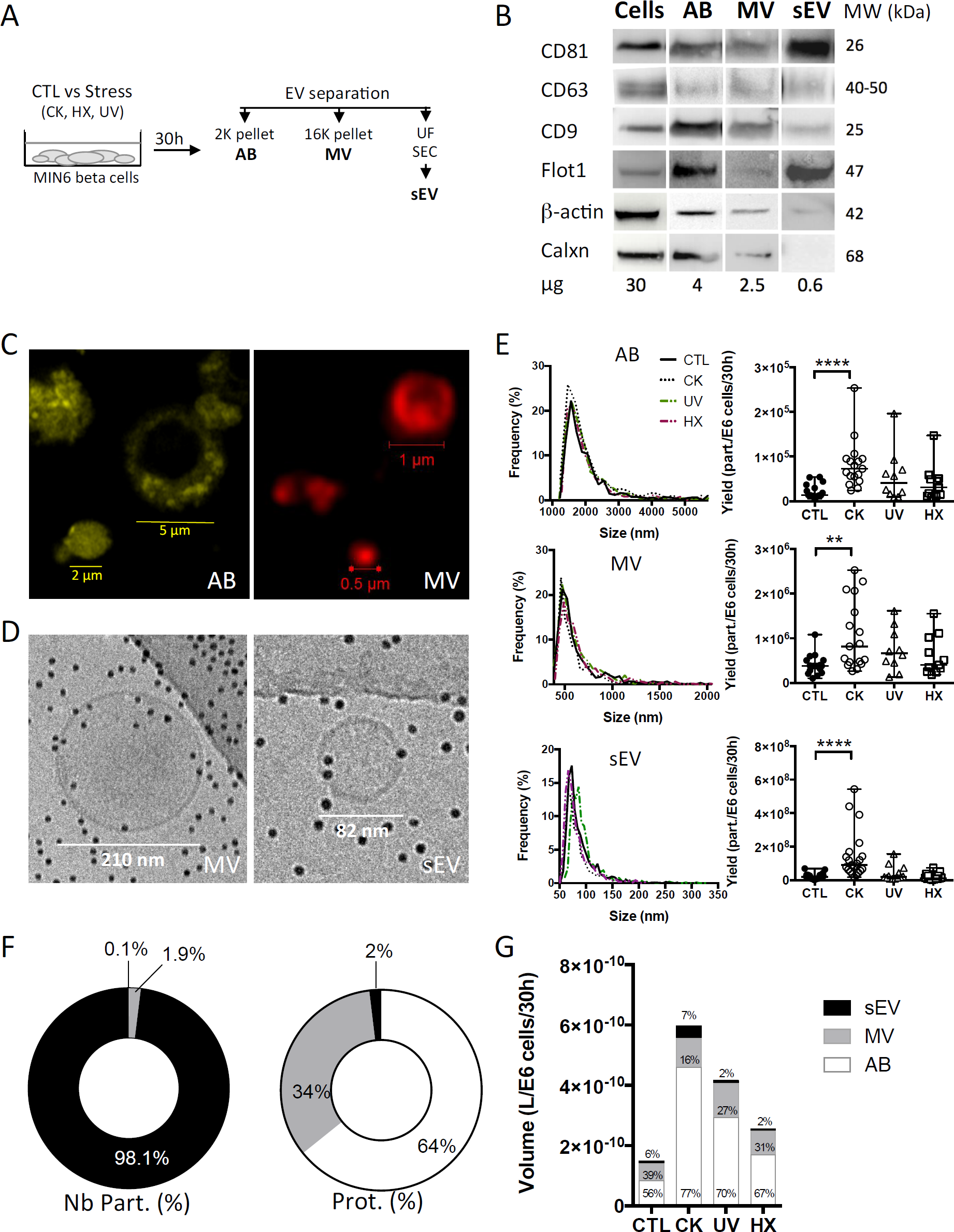
Apoptotic beta cells release a heterogeneous population of EV. (A) After exposure to stress, MIN6 cells were cultured for 30h prior to EV separation by differential centrifugation, ultrafiltration (UF) and size-exclusion chromatography (SEC). (B) Representative Western blot images of markers of extracellular vesicles (membrane CD81, CD63, CD9, Flot1, cytosolic β-actin) or cells (Calxn). (C) Confocal images of EV from untreated MIN6 cells stained with MB-Cy3 (AB) or MB-Cy5 (MV) (D) Cryo-electron microscopy images of MV and sEV. (E-G) The particle size distribution of EV subpopulations was determined by TRPS analysis. Histograms show the particle size distribution of samples from one representative experiment. Scatter dot plots represent the number of particles recovered per million of cells (data from n=10-21 independent experiments; median with range). Yields of EV obtained from MIN6 cells after treatment were compared to the yield from untreated controls using the Kruskal-Wallis test (*\*\*P*<0.01 *\*\*\*\*P*<0.0001). (F-G) Relative quantities of material released inside EV based on TRPS and Bradford protein content analyses of extracellular vesicles harvested per million of MIN6 cells after 30h of culture. Relative volumes occupied by vesicle subtypes are estimated based on mean sizes and concentrations measured by TRPS. (F) Percentage of numbers and total protein of EV subtypes derived from untreated control cells. Results are depicted as median percentages from n=7-9 replicates from independent experiments. (G) Volumes (median) measured for EV from treated and control MIN6 cells. Results from n=10-17 independent experiments are shown.

Staining with the lipid probe MemBright, recently developed by Collot and colleagues ^46,47^, showed a heterogeneous population of round-shaped vesicles in the AB and MV fractions (Fig.2C). Cryo-electron microscopy images of MV and sEV clearly ascertained the presence of a lipid bilayer membrane surrounding the vesicles (Fig.2D, Suppl.Fig.2). AB exceed the upper size limit of cryo-electron tomography (1µm) and have therefore not been analysed using this technique. TRPS analysis of the EV showed a mode size (median (range)) of 1548 (1368-1790) nm, 510 (455-563) nm, and 76 (61-120) nm for AB, MV, and sEV, respectively. None of the treatments had a significant effect on the mode or mean size of the vesicles (Fig.2E and Table 2). In healthy beta cells, large EV (AB and MV) represent 93% of the volume and 98% of the protein content of the vesicles all together, but less than 2% of the number of particles (Fig.2F). In pro-inflammatory conditions, the secretion of AB, MV and sEV was significantly enhanced as shown by the 5.5-fold, 2.1 and 4.5-fold increases of the number of particles recovered per million of cells, respectively. Commensurate to the rise in the number of vesicles, the volume occupied by the CK-EV (all subtypes) increased 4.0-fold against 2.8-fold for UV and 1.7-fold for HX EV (Fig.2G, Table 1). In line with the particle size, the number of particles per microgram of protein is much higher in MV (> 25 times) and sEV (>1E4 times) than in AB. As expected, this ratio remained constant in conditions of stress for MV and sEV, but curiously increased in AB derived under conditions of stress (CK vs CTL; p= 0.0084). In the absence of noticeable changes in the particle’s volumes measured by TRPS, this increase hints to changes in the vesicle’s protein content. Cytoplasmic vacuolation and inclusion of organelles and DNA fragments during the process of apoptosis possibly reduced the proportion of proteins in AB from CK-treated in comparison to AB from untreated cells.

**Table 1.**
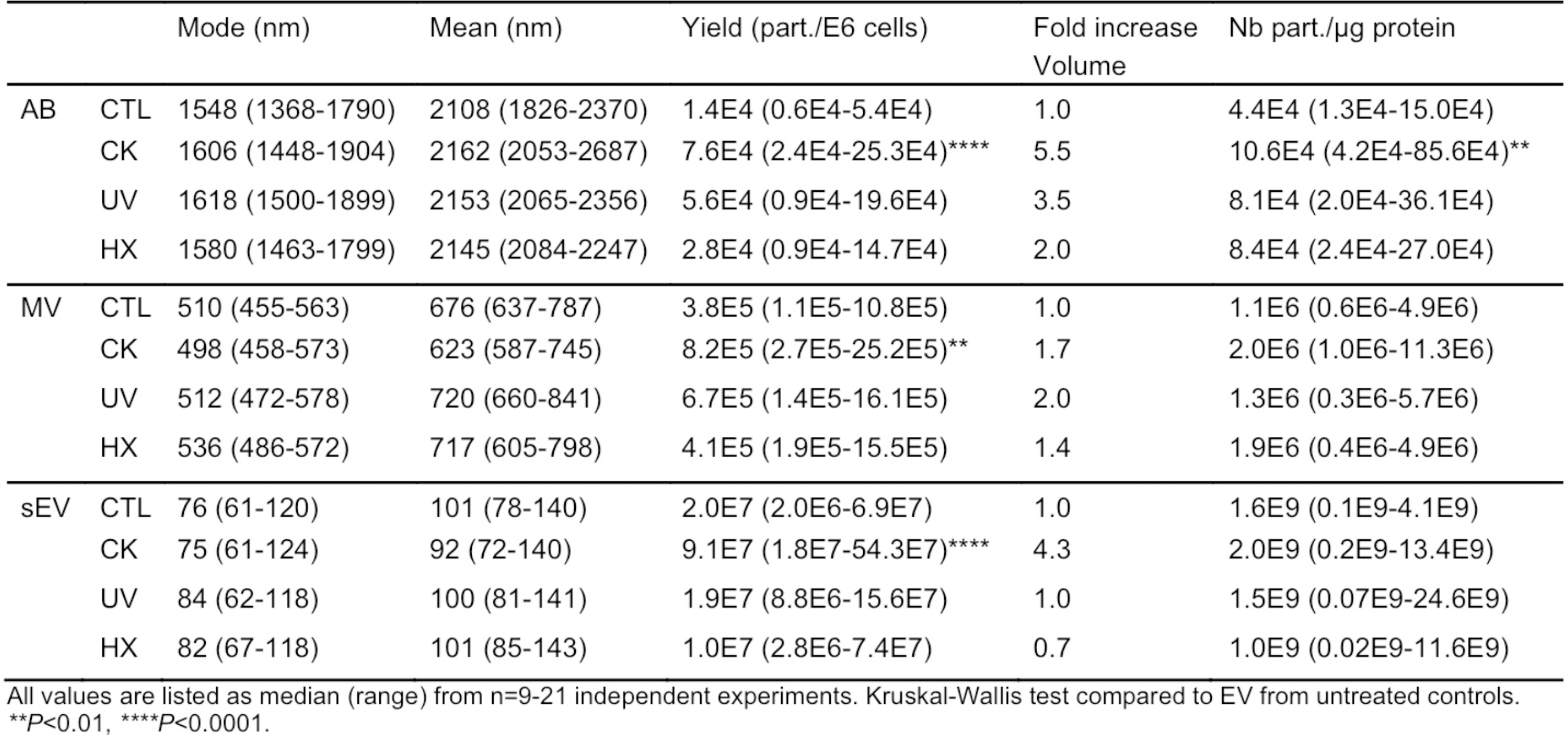
Physical properties of beta EV.

Earlier studies provided evidence that human and mouse beta EV contain major auto-antigens of type 1 diabetes such as GAD65, islet-associated antigen 2, Znt8, and insulin ^12-15^. Out of these, insulin is the most prominent islet autoantigen, highly abundant in beta cells and was used here to ease monitoring of autoantigen partition. To explore how autoantigens partition into beta EV in normal and pathological conditions, we quantified the amount of total insulin comprising pro-insulin and mature insulin in EV subpopulations by ELISA (Fig.3A). Data obtained on EV from healthy cells showed that a majority of vesicle-associated insulin was exported inside large vesicles (Fig.3B). The absolute insulin content (median (range)) was in the range of 7 × 10^−14^ (1 × 10^−14^ – 1 × 10^−12^) g/part. in AB, 1 × 10^−15^ (1 × 10^−16^ – 7 × 10^−15^) g/part. in MV down to 8 × 10^−19^ (2 × 10^−19^ – 4 × 10^−18^) g/part. in sEV, in line with the volume of the particles (Suppl.Fig.3A). In comparison to AB, a 1.5-fold and 4.6-fold lower median insulin concentration was measured respectively inside MV and sEV, where insulin represented 0.1 (0.0-0.3)% of the vesicles’ protein content (Fig.3C and Suppl.Fig.3B).

**Fig3.**
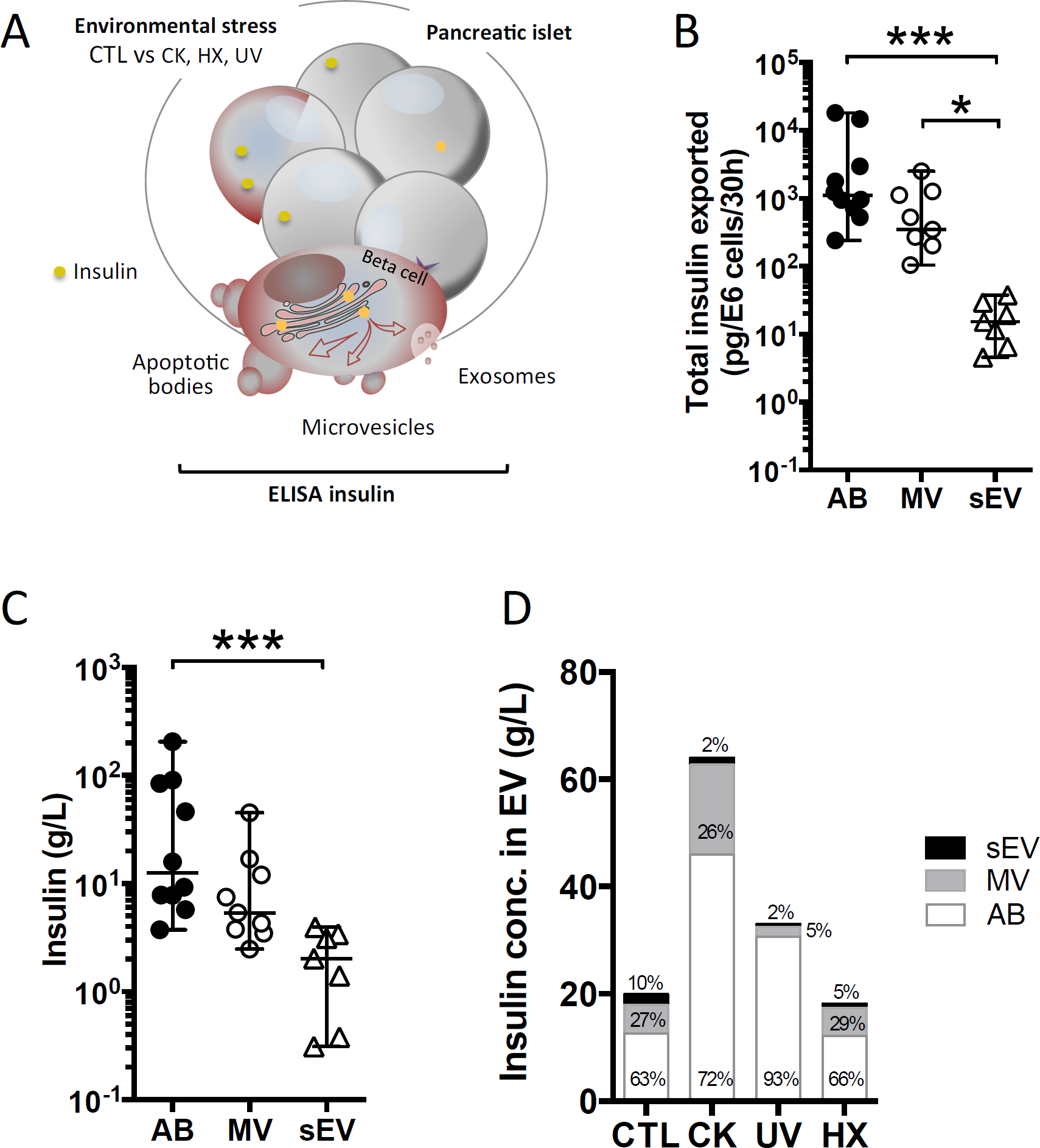
Differential partition of the autoantigen insulin inside EV. (A-D) After 30h of culture, EV were collected from supernatants from CTL, CK, UV or HX MIN6 cells and assessed for total insulin (pro-insulin and mature insulin) content by ELISA. (B-C) On EV derived from untreated control cells (CTL) (B) Quantities of insulin measured inside AB, MV and sEV collected per million of CTL cells. (C) Insulin concentrations measured in association with AB, MV and sEV from CTL MIN6 cells. (D) Evolution of sorting of insulin towards EV following exposure to stress. Data from n=7-11 independent experiments are shown with median and range and compared using the Kruskal-Wallis test (*\*P*<0.05 *\*\*\*P*<0.001).

Following exposure to stress, insulin export was markedly enhanced in AB and sEV derived from CK-treated cells, whereas no significant changes were perceived in EV derived from cells cultured exposed to hypoxia or UV-irradiation (Fig.3D and Suppl.Fig.3, Suppl.Table1). This increase in insulin export relied on enhanced EV export, as the concentration of insulin inside the EV subtypes did not change following treatment.

MiRNA may act as adjuvants in immune activation and recently six miRNA with the potential to bind directly to the TLR-7 receptor of innate immunity have been described in MV and sEV ^23,24,26-28,48,49^. Here we wanted to investigate how TLR-binding miRNA are sorted into EV and whether their expression in these vesicles changes in situations of stress.

With the aim to compare TLR-binding miRNA expression in an equal amount of cells and large and small EV derived thereof, a synthetic ath-miR-159a was spiked into all samples prior to RNA extraction. After RT-qPCR amplification, relative quantities in samples were normalised with respect to this exogenous control as well as cells or EV derived from untreated control cells (Fig.4). The results obtained show a significant up to 3-fold drop in the expression of TLR-binding miRNA in cells in situations of pro-inflammatory stress, in parallel to an up to 13- and 48-fold increase in LEV and sEV, respectively. Although less pronounced, similar trends were observed in cells and LEV obtained under genotoxic and hypoxic conditions. No changes of TLR-binding miRNA expression were observed in sEV under genotoxic and hypoxic conditions. This enhanced TLR-binding miRNA release in vesicles could be explained either by an increase in the release in EV or by a higher concentration of these miRNA inside the vesicles. To answer this question, the RT-qPCR data was further normalised to the number of particles present in the sample as determined by TRPS. The results presented in Suppl.Fig.4, revealed an evened out expression of TLR-binding miRNA in LEV in all situations. In contrast, four to five -fold higher quantities of all TLR-binding miRNA except for miR-29b were detected in sEV secreted under proinflammatory conditions. Taken together, these data suggest that beta cell stress and in particular, the exposure of beta cells to cytokines, favours export of immune stimulatory miRNA into EV and enrichment in sEV.

**Fig4.**
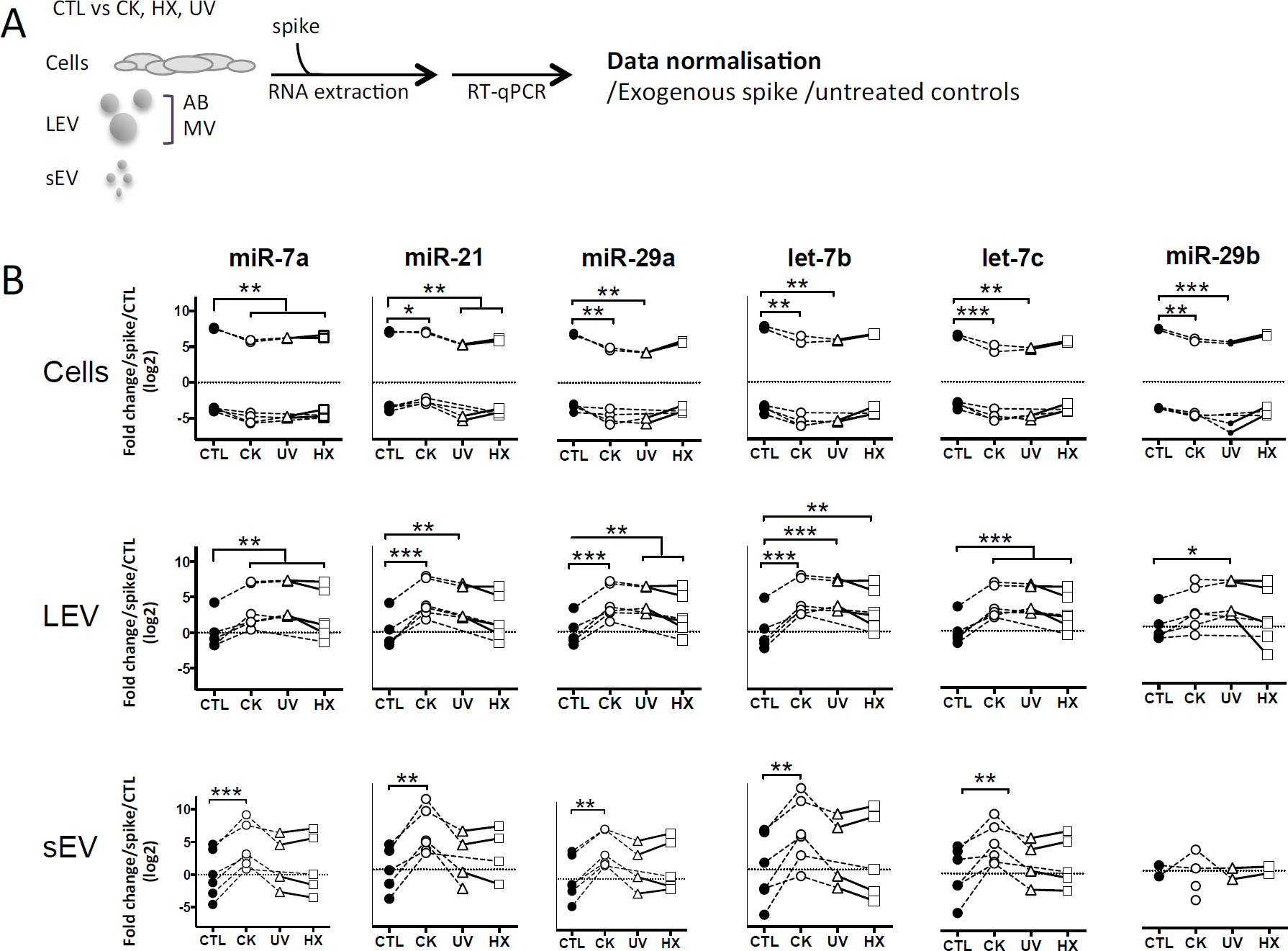
Beta cell stress favours export of TLR-binding miRNA in EV. (A) Following exposure to stress, MIN6 cells were cultured for 30h followed by isolation of large EV (comprising AB and MV) and sEV. All samples were spiked with an exogenous control prior to total RNA extraction and processed for quantitative RT-PCR. After amplification, relative quantities were normalised with respect to the spike and untreated controls. (B) Quantitative RT-PCR analysis of miRNA expression in a fixed number of cells and EV derived thereof. Individual replicates from 4-6 independent EV isolations are represented as fold-changes compared to untreated controls. Tukey’s range test *\*P*<0.05, *\*\*P*<0.01 and *\*\*\*P*<0.001.

Cytokines are well-known soluble mediators in cell-to-cell communication, however evidence exist that an important fraction of biologically active cytokines are released from tissues in association with EV either bound to the surface or encapsulated in the lumen of the vesicles ^50^. Owing to cytokine treatments performed in our EV production workflow, we performed a mouse 13-plex CBA to assess for interferon (IFNβ, IFNγ), interleukin (IL-1α, IL-1β, IL6, IL-10, IL-12p70, IL-17A, IL-23, IL-27), granulocytes macrophage colony stimulating factor (GM-CSF), TNFα and monocyte chemoattractant protein-1 (MCP-1; also called chemokine (C-C motif) ligand 2 (CCL2)) cytokine expression in the different subpopulations of beta EV. Prior to analysis, the EV were incubated in 0.5% Triton X-100 for 30 minutes and sonicated for 1 min. to assess for internal as well as surface-associated cytokines. Six cytokines were detected in subpopulations of MIN6 beta EV (Fig.5 and Suppl. Table1). Seven cytokines were below detection levels in all vesicles. None of the EV secreted by untreated controls exhibited the exogenous cytokines TNFα, IFNγ or IL-1β (Suppl. Table1). Trace amounts of MCP-1 and IL-23 were detected in CTL-AB and CTL-MV. Following exposure to picomolar concentrations of cytokines in the initial culture medium, MIN6 release femtograms of IFNγ, TNFα and IL-1β per million of cells in AB and, to a lesser extent, in MV. IFNγ was not detected in sEV and quantities close to detection thresholds of IL-1β and TNFα were detected in only 1 out of four and 3 out of four samples, respectively. Interestingly, MIN6 cytokine treatment stimulated a dramatic increase in the expression (median (range)) of MCP-1 in AB [358 (281– 415.5) fg/E6], MV [127.5 (65.1 – 208.8) fg/E6 cells] and sEV [16.4 (3.6 – 25.4) fg/E6 cells] and of IL-27 in AB [24.1 (15.7 – 33.4) fg/E6 cells]. Cytokine profiles of EV derived from MIN6 cells exposed to hypoxia or UV-irradiation were similar to profiles from untreated cells (data not shown).

**Fig5.**
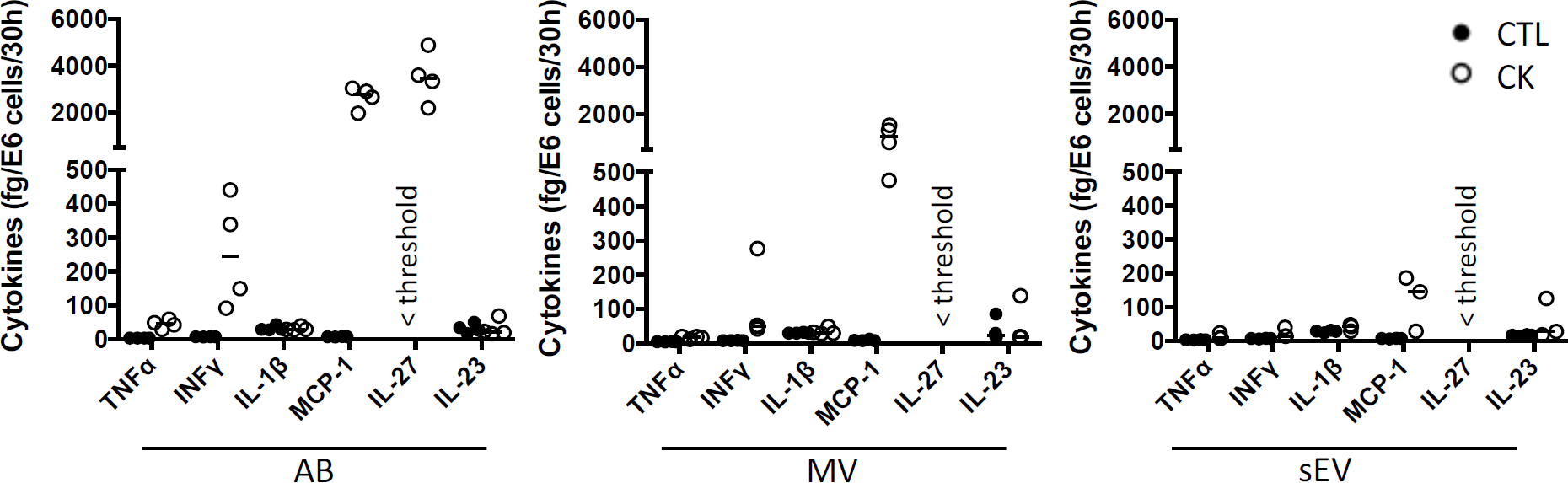
Cytokine profile of beta EV. Multiplex CBA monitoring of cytokine expression in association with EV collected from 30 hours culture supernatants from cytokine-treated MIN6 cells and untreated controls. Data (with median) from 3-4 independent experiments are shown.

APC such as dendritic cells and macrophages have been identified as recipient cells for beta EV uptake *in vitro* and *in vivo*^12,13,51,52^. For side-by-side comparisons of the potential of EV subtypes to modulate APC function, primary NOD bmDC were exposed to MIN6-derived beta EV for eighteen hours followed by flow cytometry analysis of the expression of MHC-II and co-stimulatory CD86 and CD40 molecules (Fig.6). AB derived from cytokine-treated beta cells induced a modest albeit significant up-regulation of CD40 and MHC II expression. CD86 expression remained unchanged despite a wider variation of expression. None of the MV and sEV modulated the expression of co-stimulatory molecules in bmDC (Fig.6A and data not shown).

**Fig6.**
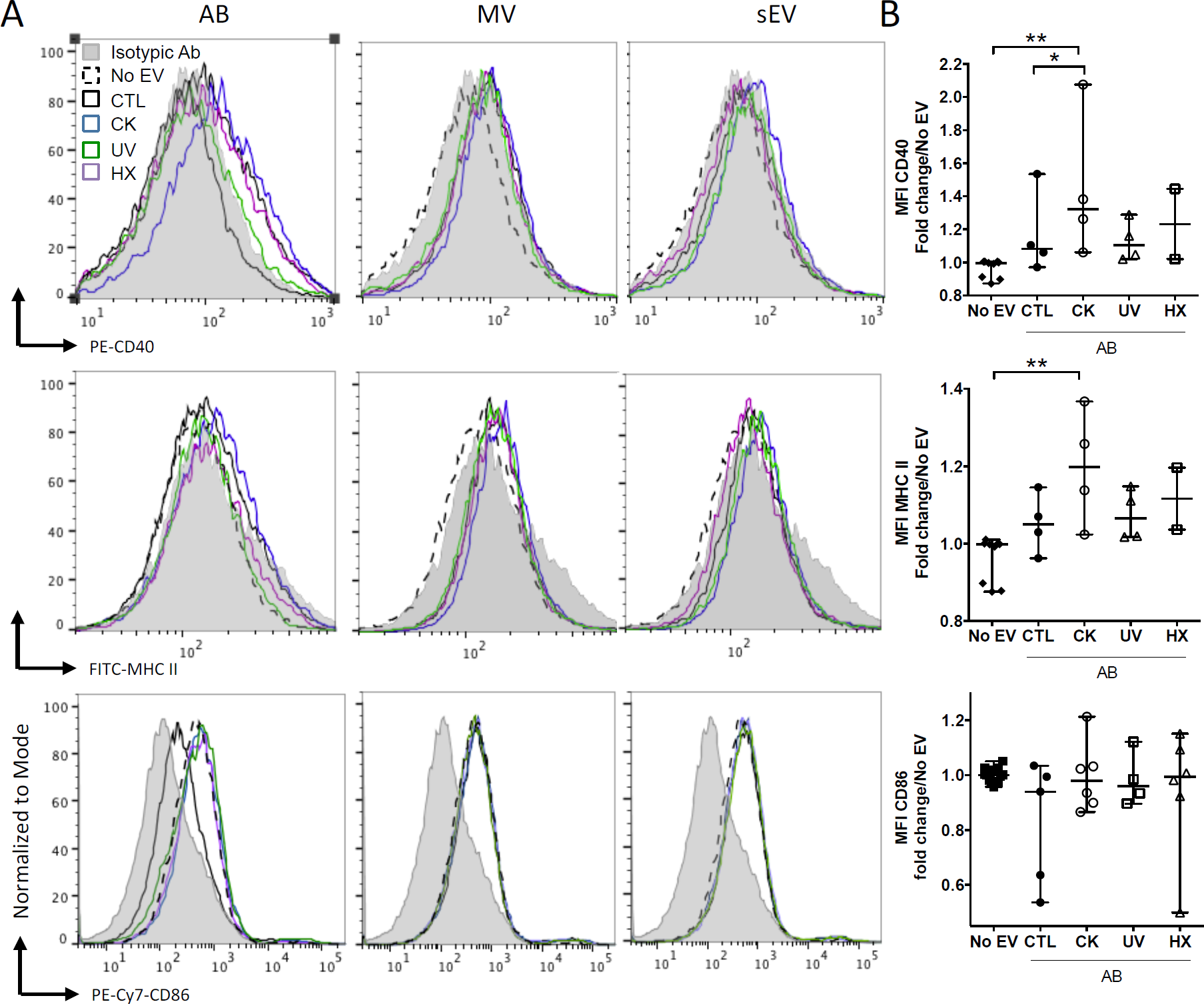
Modulation of co-stimulatory molecules in NOD bmDC by beta AB. NOD_Shi_ mice derived bmDC were incubated for 18h with AB, MV or sEV derived from 8 × 10^7^ MIN6 cells. (A) Flow cytometric analysis of the expression of the CD40, MHC II and CD86 activation markers. Histograms with dashed lines represent bmDC treated with AB derived from cytokine beta cells. Histograms with solid lines represent bmDC treated with AB from untreated beta cells. Gray histograms represent isotypic controls. Data from one representative out of 2-5 independent experiments are shown. (B) Scatter plots show the median fluorescence intensity (MFI) of co-stimulatory molecules expression in bmDC after AB treatment normalised to No EV controls (median with range). Tukey’s range test *\*P*<0.05 and *\*\*P*<0.01.

The concentration of cytokines in supernatants of murine bmDC and RAW264.7 macrophages exposed to EV in culture was assessed by CBA and ELISA (Fig.7). In bmDC, no significant influence of the EV treatments was observed on the concentrations of IL-1α, IL-1β, IL-6, IL-10, IL-12p70 or IL-23 whose levels remained low in culture supernatants, close to detection thresholds (Fig.7A and data not shown). Among cytokines expressed in beta EV derived under inflammatory situations, MCP-1 was detected in bmDC cultures with AB and MV derived from MIN6 cells cultured under inflammatory conditions (Fig.7B). MCP-1 concentrations measured in culture supernatants are 2-4 times lower than concentrations calculated for AB input into these culture, supporting the idea of an essentially passive carry-over of MCP-1. In contrast, IL-27, which was highly expressed in CK-AB, was below detection thresholds in bmDC culture supernatants (data not shown) suggesting differential kinetics of MCP-1 and IL-27 uptake, recycling or activity in bmDC. Alternatively, sustained expression of MCP-1 in culture supernatants could be explained by *de novo* cytokine production by bmDC. All AB (except UV-AB) and sEV derived under inflammatory conditions led to increased levels of TNFα in bmDC culture supernatants, superior to TNFα amounts provided by these vesicles. Though CK-MV expressed 2.5-fold higher levels of TNFα than CK-sEV, no differences in the concentration of TNFα was observed in bmDC supernatants in the presence of CK-MV in comparison to CTL-MV (Fig.7C). Co-incubation of murine RAW264.7 macrophages with AB, MV as well as sEV led to increased TNFα supernatant concentrations for EV obtained from cytokine-treated MIN6 cells, the amounts of which cannot solely be explained by passive carry-over of EV-associated TNFα. For EV from beta cells in hypoxia, AB were also able to significantly enhance TNFα secretion (Fig.7D; p=0.0122). For all EV from UV- and HX-stressed beta cells, a tendency of TNFα induction was visible. As inflammatory stress was the most potent inducer of EV release in our hands (Fig.2), the more pronounced immune effects might be caused by a higher ratio of EV to target immune cells or a minimal concentration of EV necessary for immune activation. Taken together, our results reveal modest direct or indirect activation of dendritic cells and macrophages by beta EV.

**Fig7.**
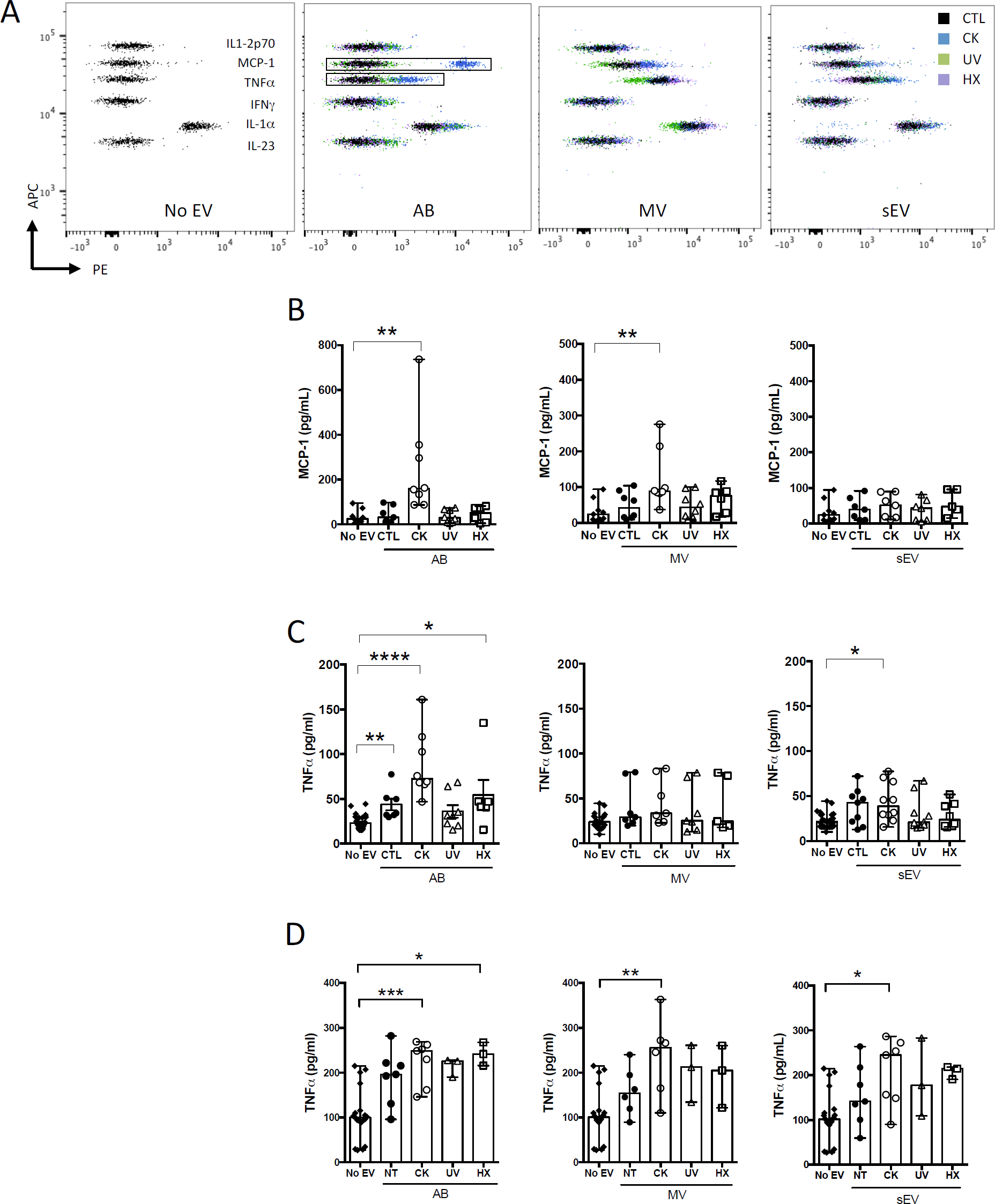
Modulation of cytokine secretion profiles in innate immune cells by beta EV. (A-C) NOD_Shi_ mice derived bmDC were incubated for 18h with AB, MV or sEV derived from 8 × 10^7^ MIN6 cells (A-B) CBA of cytokine concentration in bmDC culture supernatants. (A) Dot plots of fluorescence intensity of beads from one representative experiment are shown. (B) CBA of MCP-1 in culture supernatants from bmDC treated with AB or No EV controls. Data from 6-7 independent experiments are shown (median with range). (C) TNFα supernatant levels as confirmed by ELISA. Data from 6-8 independent experiments are shown. (D) RAW264.7 macrophages were treated for 18h with AB, MV or sEV derived from 8 × 10^7^ MIN6 cells. TNFα cytokine levels in culture supernatants were measured by ELISA. Data from 3-6 independent experiments are shown. Kruskal Wallis test *\*P*<0.05, *\*\*P*<0.01 and *\*\*\*P*<0.001.

## Discussion

Living cells release heterogeneous populations of EV that constitute a means for surrounding and distant tissue crosstalk. Beta sEV have been shown to drive innate and adaptive prodiabetogenic immune responses, but the functional diversity of the beta secretome as a whole, and the impact of beta cell stress on the beta EV repertoire have not been explored yet.

We observe that experimental exposure of MIN6 beta cells to either inflammatory cytokines, low oxygen tension or UV-irradiation rapidly induces ER-stress and subsequently apoptosis. While p-eIF-2a, indicative of translational attenuation is observed in all situations of stress, the apoptosis-mediating transcription factor CHOP is barely detected in UV-treated MIN6 cells. This is in line with earlier observations that DNA damage through UV irradiation alone is insufficient to induce CHOP expression ^53^. Although low doses of cytokines were used here in comparison to similar beta cell studies ^17,54,55^, the percentage of caspase-3/7 positive beta cells were 4-fold higher in CK-treated cells than in cells following UV irradiation or cultured under hypoxia, illustrating the particular propensity of beta-cells to undergo apoptosis in response to inflammatory stressors ^56^.

Stress and inflammation have been repeatedly reported to enhance EV secretion, including from beta cells ^12,57^. Here, quantitative side-by-side comparisons of EV subtypes isolated from an equal amount of beta cells, reveal a consistently higher release of EV exclusively under inflammatory conditions. To what extent, the five-, two- and four-fold increases in the number of particles observed for AB, MV and sEV, respectively, impact downstream immune responses presumably depends on the nature of their cargo.

Indeed, in response to stress, cancer cells secrete EV which have been shown to contribute to survival of surrounding cancer cells and drug resistance ^58^. In contrast, sEV from cytokine-treated beta cells induce beta cell apoptosis in naïve beta cells ^17^ suggesting an EV-mediated spread of inflammatory cellular constituents. Following cytokine exposure, it has been shown that chaperones of the UPR promoting DAMP-signalling, namely calreticulin, Gp96, ORP150 and the heat-shock protein HSP-90α are packed into beta sEV ^12,54^. Beta EV have also attracted interest for their aptitude to transport self-antigens ^12-15^. In the present study, the partition of the highly abundant insulin protein was monitored in EV subpopulations derived under normal and pathological conditions. In our hands, 1.5% of untreated MIN6 cells continuously undergo apoptosis in culture. These apoptotic cells release AB containing 91% of the particulate secretome’s insulin content in line with the role of AB in the disposal of cellular material in efferocytosis ^59-62^. Exposure to inflammatory triggers up-regulates not solely the number of vesicles released, but also the absolute amount of insulin exported with significantly higher levels of insulin measured in association with EV produced by cytokine-treated cells. Our results converge with others obtained for TLR-binding miRNA expression inside large and small EV. Inflammatory cytokines and to a lesser extent hypoxia and UV-irradiation promote TLR-binding miRNA efflux from the cell. Interestingly, this increase relies on the enhanced secretion of LEV with an unchanged miRNA content in contrast to enhanced secretion paralleled by a rise in the relative quantities of these miRNA sequences packed inside individual sEV. Taken together, our data provide strong evidence that sorting of immunogenic material inside subpopulations of EV is not a random process and is profoundly altered in inflammatory settings.

MIN6 beta EV in the steady state contain low levels of MCP-1 and IL-23 and undetectable levels of a panel of eleven other cytokines involved in pathways of inflammation. Experimental exposure of MIN6 cells to inflammatory cytokines engenders drastic changes in the expression of the same cytokines in large AB and MV, but also of *de novo* produced MCP-1 (in AB, MV, and to a lower extent in sEV) and IL-27 (AB) cytokines. Passive exogenous cytokine carry-over is obviously a concern in the interpretation of immune functions of EV. However, it has to be stressed that beta cells facing immune insults *in situ*, most likely discard cytokines of beta or immune cell origin in an analogous manner that is to say inside large rather than small vesicles according to our data.

Several studies, including genome-wide association studies ^63-65^, demonstrate pathogenic roles of the cytokine IL-27 in T1D. Transgenic NOD IL-27 receptor knockout mice are resistant to disease and blockade of IL-27 delays T1D onset in NOD mice^66,67^. MCP-1 is a chemokine involved in immune cell recruitment. Exported in exosomes, MCP-1 has been shown to contribute to inflammation in nephropathies ^68,69^. In the context of T1D, chemotaxis assays showed that subnanomolar amounts of MCP-1 produced by beta cells are sufficient to attract monocytes ^70^. It has been shown earlier, that mouse and human islet cells constitutively express MCP-1 and produce high levels of MCP-1 peaking at 6h of incubation in response to proinflammatory cytokines ^71,72^. In islet transplantation, MCP-1 is inversely correlated to islet graft function ^70,73^ and attempts to block MCP-1 signalling successfully improve graft survival^74^. Furthermore, T-lymphocyte exosomes induce MCP-1 expression and apoptosis in beta cells^75^ illustrating the importance of MCP-1 in beta cell inflammation and failure. MCP-1 stimulation on its own results in aberrant sorting of immune regulatory miRNA into extracellular vesicles^76^ in line with observations made on TLR-binding miRNA in our study. It is thus conceivable that molecular mediators of inflammation as chemokines and immunostimulatory miRNA establish and mutually maintain inflammation.

Added to culture of bmDC derived from diabetes-prone NOD mice, EV from cytokine-treated beta cells up-regulate moderately the surface expression of MHC class II and co-stimulatory CD40 molecules. In these experiments, an EV donor to recipient ratio of 1:200 was used, which would be equivalent to 5 DC in an averaged sized islet containing 1000 beta cells, a plausible proportion in the inflamed pancreas at disease initiation. EV from CK-treated MIN6 cells exert the strongest immune effects, which could be due to cumulative effects of cargo quantity (autoantigens, proinflammatory miRNA, endogenous cytokines), increased EV release and cytokine carry-over. At least two facts argue against cytokine carry-over as the only responsible for the observed immune effects. First, AB derived under hypoxia devoid of EV-associated cytokines also significantly induced TNFα secretion in RAW264.7 macrophages. Second, IL-27 highly expressed in CK-AB was not detected in bmDC culture supernatants. Lastly, we showed earlier that TNFα secretion in RAW264.7 macrophages induced by sEV derived from untreated MIN6 cells is positively correlated to the amount of particles in culture ^26,51^.

The relevance of these quantitative and qualitative differences of subsets of apoptotic beta EV have to be weighted with regard to the interplay of these vesicles with cellular effectors of immunity *in vivo*. Obviously, enhanced EV release in situations of stress engenders higher EV to immune cell ratios. This fact should be considered in EV biological activity assays, which are frequently based on treatments with a constant number of particles. AB are known to express “find me” and “eat me” signals leading to rapid elimination by patrolling phagocytes ^77^. Conceivably, AB from stressed beta cells constitute a critical source of chemo-attractants, beta self-antigens and danger signals that could infer with the otherwise immune silent elimination of the dead by efferocytosis. In contrast, nanosized vesicles such as sEV and small MV have half-lives of minutes to hours *in vivo* ^78^. They are in the ideal size range for transport in interstitial fluids and have been shown to efficiently diffuse to secondary lymphoid organs as spleen and draining lymph nodes ^79-81^. Thereby, MV and sEV from inflamed pancreatic islets could have implications in immune regulation by aberrant autoantigen and immune-stimulatory miRNA expression at nearby as well as distant sites. Taken together, our findings highlight the profound impact of inflammation in comparison to other stressors on the beta EV repertoire. Centred on stress, the induction of markers of activation and mediators of inflammation (with the exception of IL-10) by beta EV are analysed in the present work. Further investigations are required to dissect the mechanisms of potential protective versus pathological roles of EV subspecies from healthy and stressed beta cells in T1D development.

## Supporting information

SupplFig1_Giri et al

## Acknowledgements

The authors are most grateful to Prof. J.I Miyazaki (University Medical School, Osaka, Japan) for the MIN6 cell line, to M. Collot (CNRS UMR7213, Strasbourg, France) for Membright, and to L. Duchesne (IGDR, UMR6290, Rennes) for mix-capped gold nanoparticle preparation. The authors acknowledge valuable help from B. Blanchet, O. Andrieu and M Leble for assistance with animal care, the APEX platform UMR703 INRA Oniris for confocal imaging and the Pays de la Loire & Ministry (PhD KG) & French National Research Agency (ANR-10-IBHU-005) for financial support.

## Author contributions

K.G., L.dB., D.J., J.-M.B., G.M. and S.B. designed the experiments. K.G., L.dB., M.L. and S.B. produced and characterized the EV and performed functional assays. D.J. carried out immunohistochemical analyses. R.F. and L.D. did the confocal microscopy analyses. A.D. and D.C. carried out cryo-tomography analyses. All authors contributed to data analysis. G.M. performed statistical analysis with RStudio. K.G., S.B., G.M. and J.-M.B. wrote the paper. All authors have read and approved the manuscript.

## Supplementary figures

**Suppl.Fig1.**
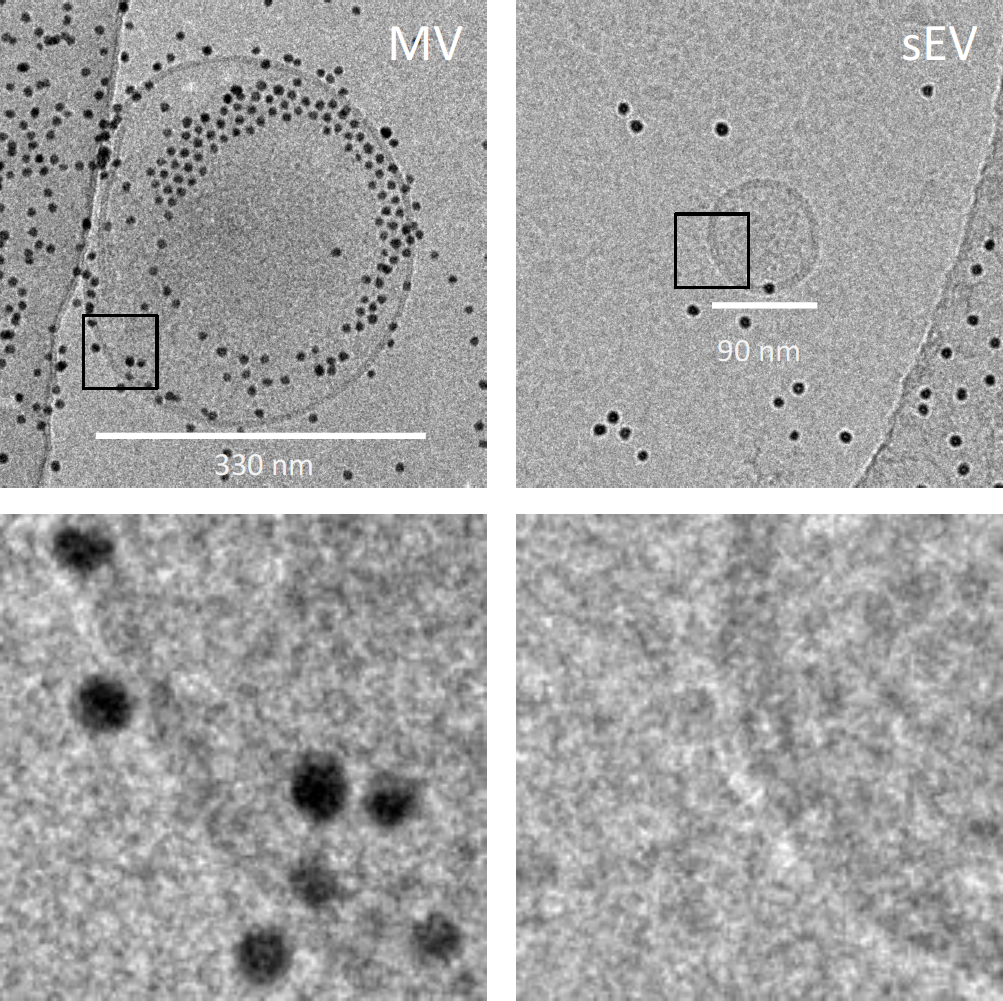
Video caspases. For monitoring of kinetics of apoptosis in live cells, a 30h time-lapse acquisition was performed on CK and untreated control MIN6 cells on a LSM780 confocal microscope (Zeiss, Oberkochen, Germany) in a 5% CO_2_, 20% O_2_, 37°C atmosphere. Apoptosis was assayed by fluorescent caspase-3/7 substrate cleavage staining. Briefly, 3×10^5^ cells/cm^2^ MIN6 cells were cultured in eight-chamber labteks with coverslips. After overnight culture, cells were switched to OptiMEM 1% FCS production medium and exposed to cytokines or left untreated and 2 mM caspase-3/7 detection reagent (Fisher Scientific) were added. Images were captured every 15 minutes.

**Suppl.Fig2.**
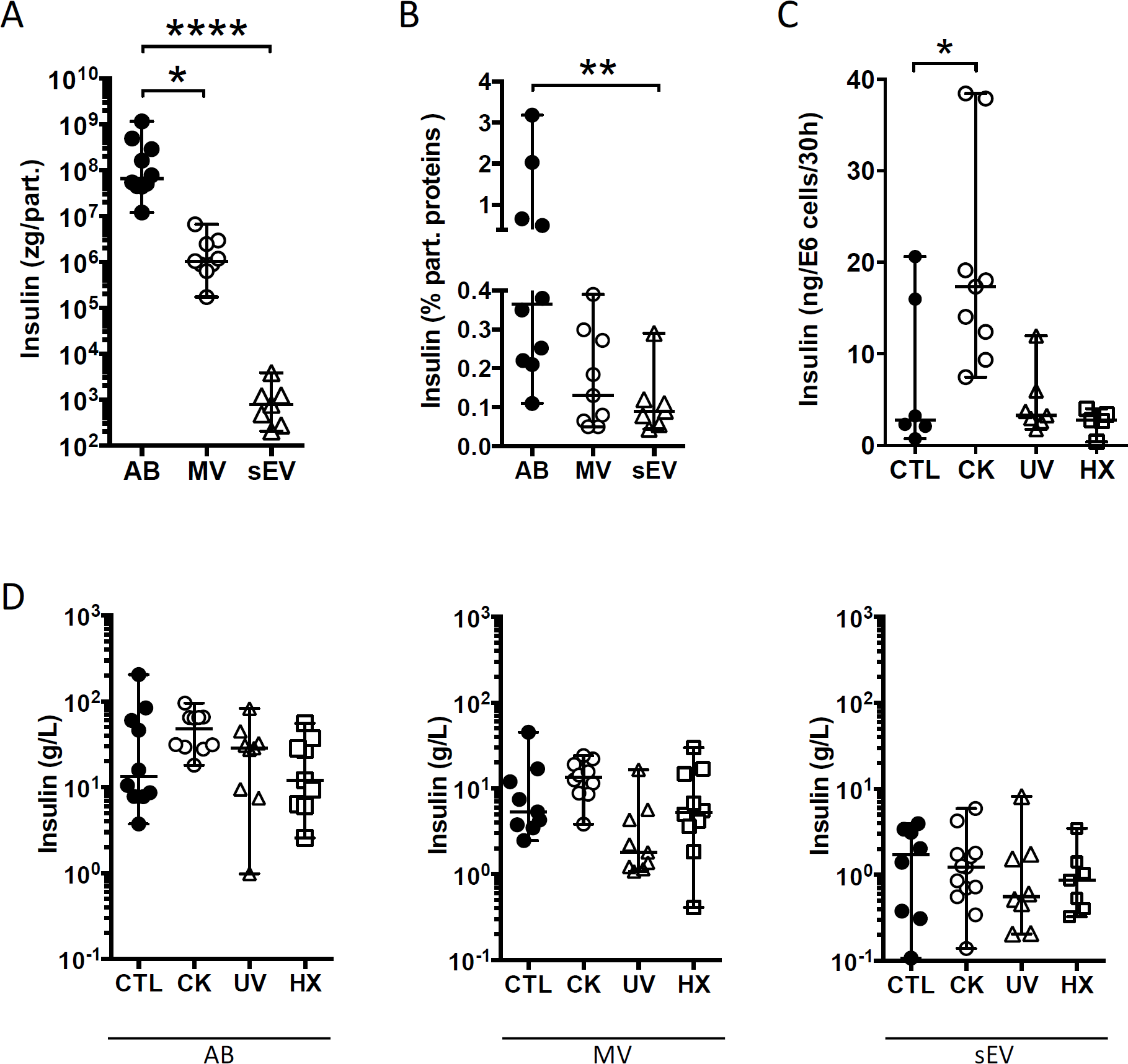
Cryo-electron microscopy of EV from untreated MIN6 cells. Images of entire vesicle of MV and sEV are represented in the top row with zooms on inserts depicted in the bottom row. Images were acquired at a nominal magnification of x 29,000.

**Suppl.Fig3.**
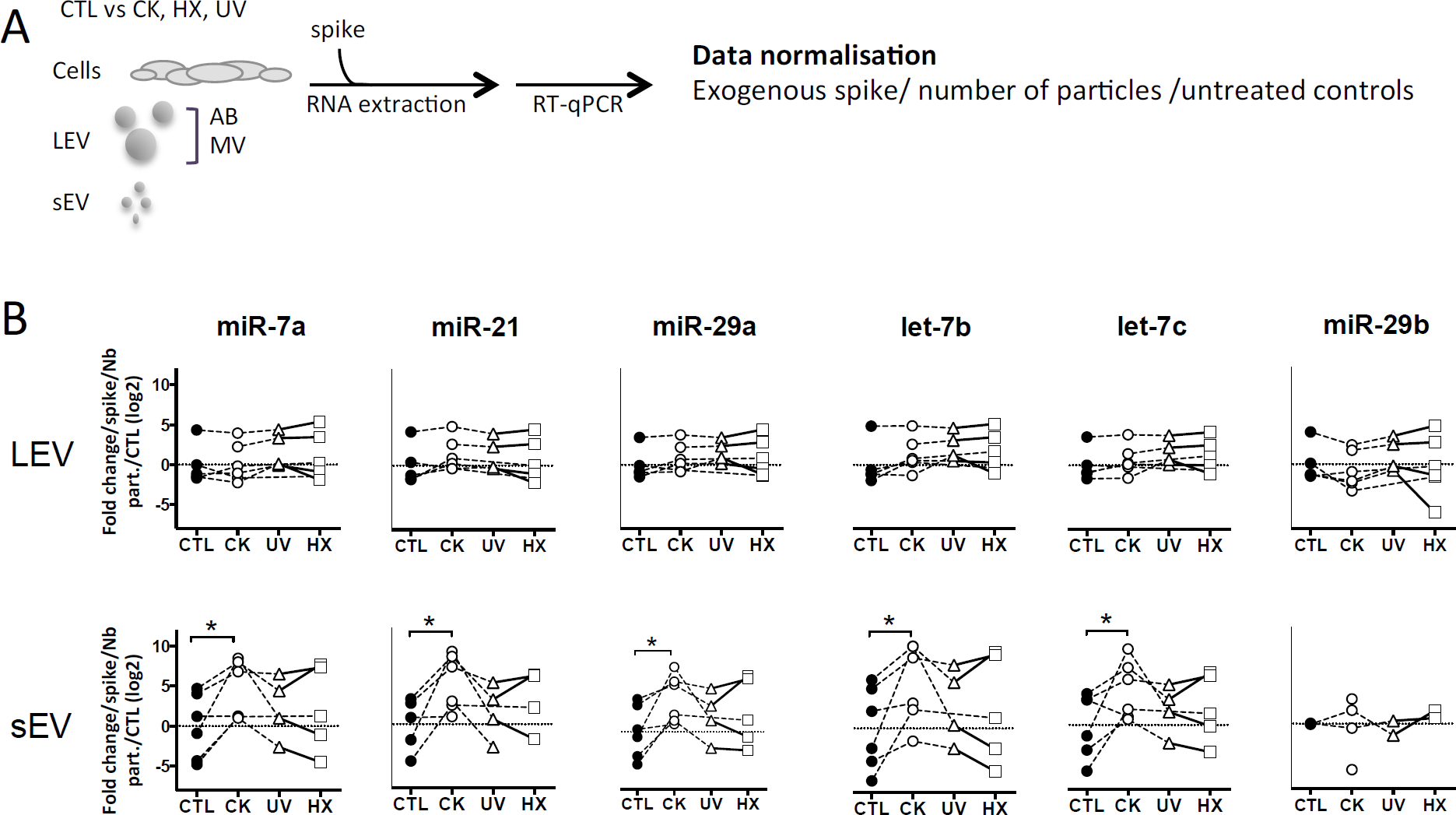
Differential partition of the autoantigen insulin inside EV. After 30h of culture, EV vesicles were collected from supernatants from MIN6 cells with (A-B) no treatment (C-D) all treatment situations and assessed for total insulin (pro-insulin and mature insulin) content by ELISA. (A) Absolute quantities of insulin measured inside AB, MV and sEV reported to the number of particles. (B) Percentage of insulin out of the total protein content in AB, MV and sEV. (C) Sum of quantities of insulin released inside AB, MV and sEV per million of producer cells. (D) Concentration of insulin inside EV. Data from n=7-11 independent experiments are shown with median and range and compared using the Kruskal-Wallis test (*\*P*<0.05 \**\*P*<0.01 *\*\*\*\*P*<0.0001).

**Suppl.Fig4.**
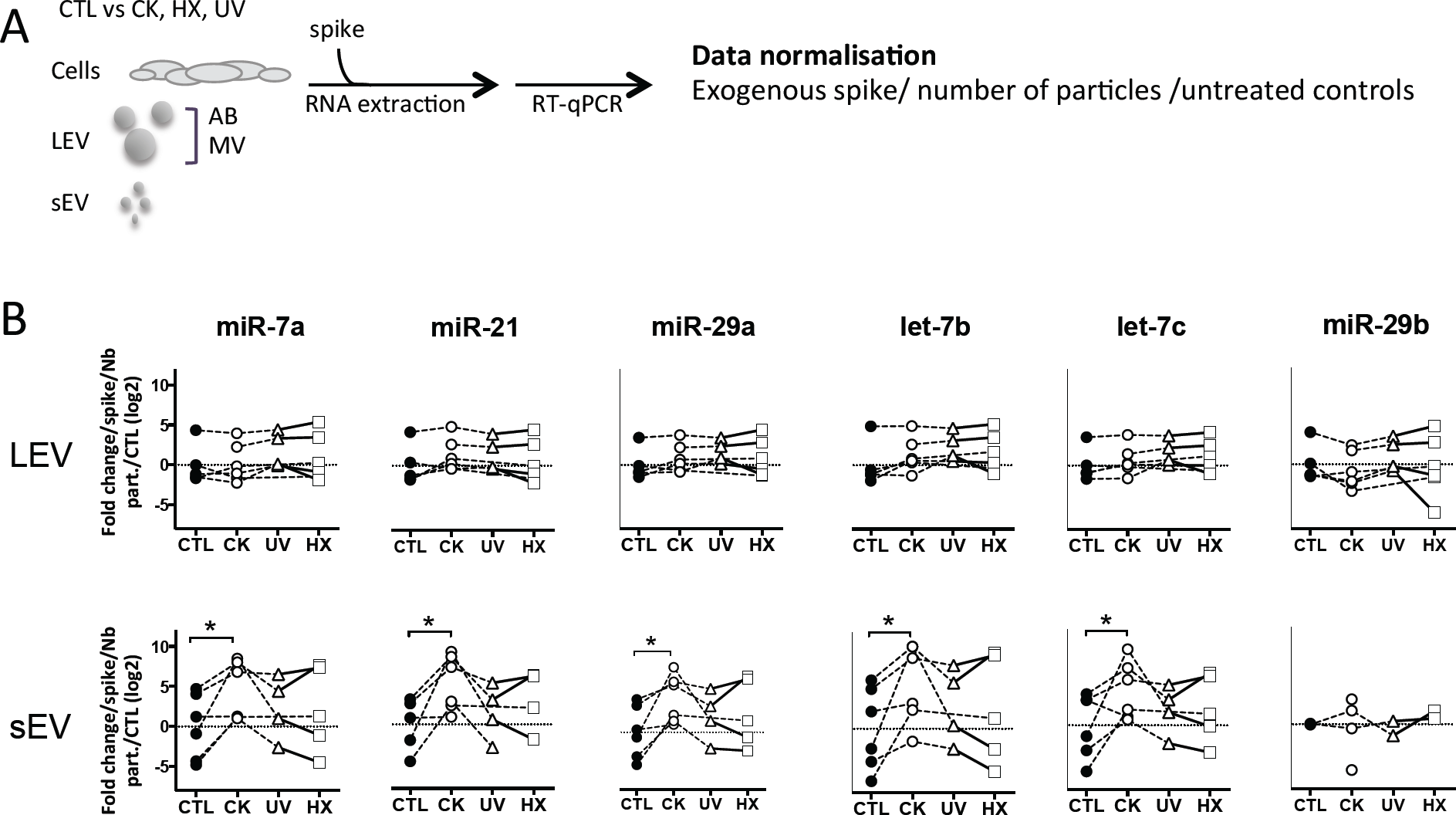
Beta cell stress favours export of TLR-binding miRNA in EV. (A) Following exposure to stress, MIN6 cells were cultured for 30h followed by isolation of large EV (comprising AB and MV) and sEV. All samples were spiked with an exogenous control prior to total RNA extraction and processed for quantitative RT-PCR. After amplification, relative quantities were normalised with respect to the spike, the number of particles as determined by TRPS analysis and untreated controls. (B) Quantitative RT-PCR analysis of miRNA expression in a fixed number of cells and EV derived thereof. Individual replicates from 4-6 independent EV isolations are represented as fold-changes compared to untreated controls. Tukey’s range test *\*P*<0.05, *\*\*P*<0.001 and *\*\*\*P*<0.001.

**Suppl. Table 1.**
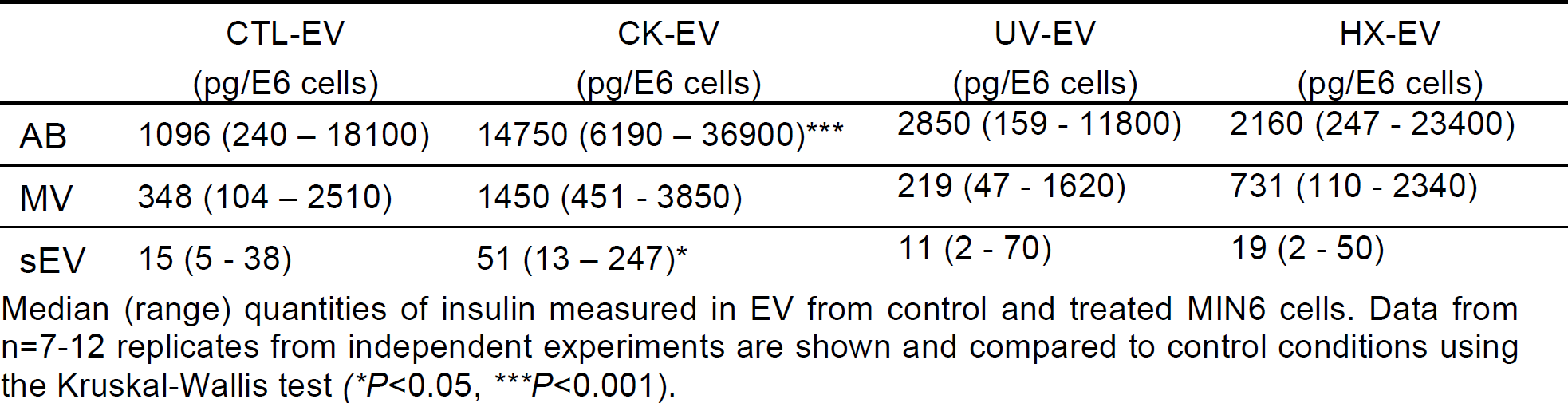
EV- associated insulin.

|  | CTL-EV<br>(pg/E6 cells) | CK-EV<br>(pg/E6 cells) | UV-EV<br>(pg/E6 cells) | HX-EV<br>(pg/E6 cells) |
| --- | --- | --- | --- | --- |
| AB | 1096 (240 – 18100) | 14750 (6190 – 36900)*** | 2850 (159 - 11800) | 2160 (247 - 23400) |
| MV | 348 (104 – 2510) | 1450 (451 - 3850) | 219 (47 - 1620) | 731 (110 - 2340) |
| sEV | 15 (5 - 38) | 51 (13 – 247)* | 11 (2 - 70) | 19 (2 - 50) |

Median (range) quantities of insulin measured in EV from control and treated MIN6 cells. Data from n=7-12 replicates from independent experiments are shown and compared to control conditions using the Kruskal-Wallis test (\* $P < 0.05$ , \*\*\* $P < 0.001$ ).

**Suppl. Table 2.**
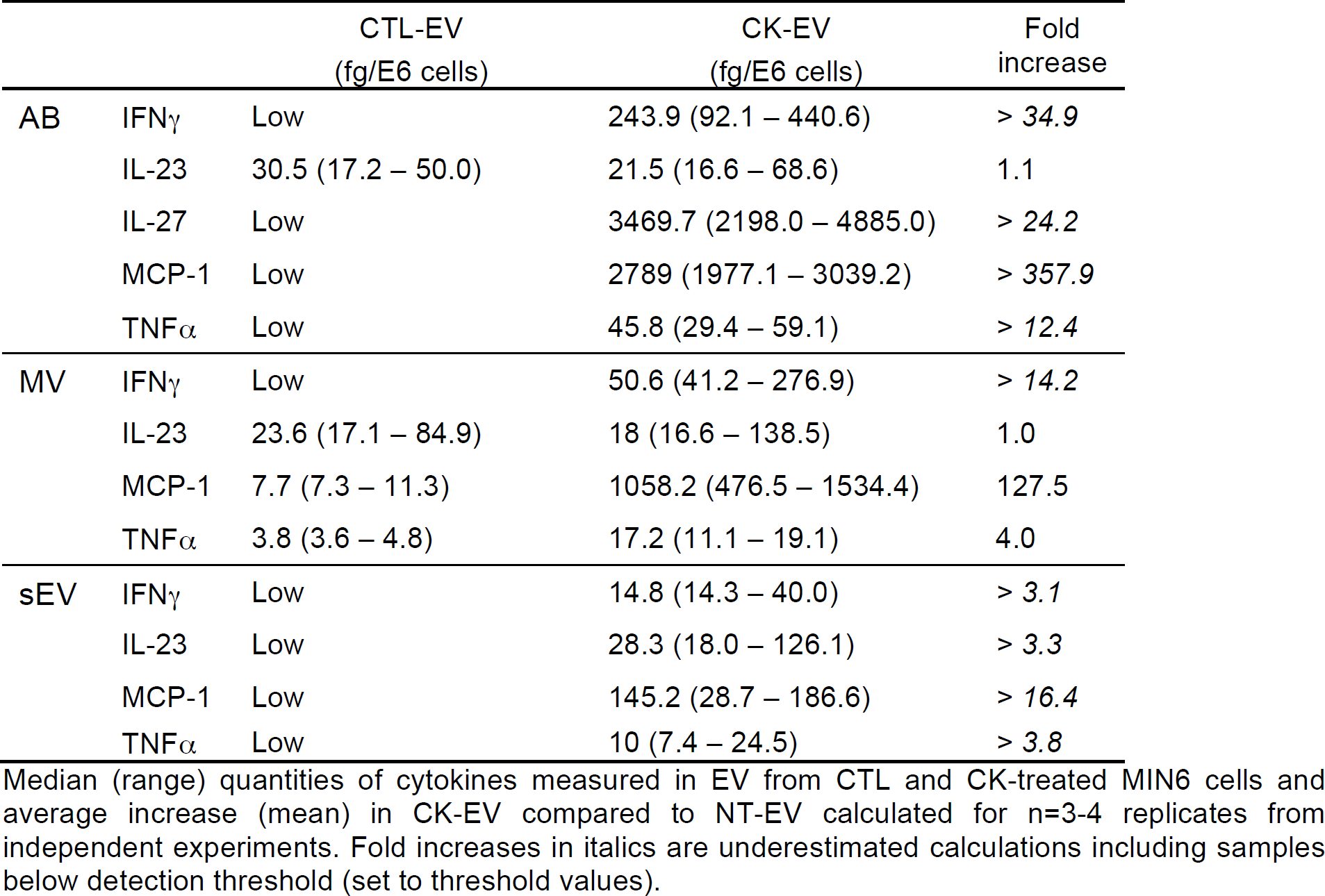
EV- associated cytokines.

|  |  | CTL-EV<br>(fg/E6 cells) | CK-EV<br>(fg/E6 cells) | Fold<br>increase |
| --- | --- | --- | --- | --- |
| AB | IFNγ | Low | 243.9 (92.1 – 440.6) | > *34.9* |
|  | IL-23 | 30.5 (17.2 – 50.0) | 21.5 (16.6 – 68.6) | 1.1 |
|  | IL-27 | Low | 3469.7 (2198.0 – 4885.0) | > *24.2* |
|  | MCP-1 | Low | 2789 (1977.1 – 3039.2) | > *357.9* |
| | TNFα | Low | 45.8 (29.4 – 59.1) | > *12.4* |
| MV | IFNγ | Low | 50.6 (41.2 – 276.9) | > *14.2* |
|  | IL-23 | 23.6 (17.1 – 84.9) | 18 (16.6 – 138.5) | 1.0 |
|  | MCP-1 | 7.7 (7.3 – 11.3) | 1058.2 (476.5 – 1534.4) | 127.5 |
| | TNFα | 3.8 (3.6 – 4.8) | 17.2 (11.1 – 19.1) | 4.0 |
| sEV | IFNγ | Low | 14.8 (14.3 – 40.0) | > *3.1* |
|  | IL-23 | Low | 28.3 (18.0 – 126.1) | > *3.3* |
|  | MCP-1 | Low | 145.2 (28.7 – 186.6) | > *16.4* |
| | TNFα | Low | 10 (7.4 – 24.5) | > *3.8* |

Median (range) quantities of cytokines measured in EV from CTL and CK-treated MIN6 cells and average increase (mean) in CK-EV compared to NT-EV calculated for n=3-4 replicates from independent experiments. Fold increases in italics are underestimated calculations including samples below detection threshold (set to threshold values).

