## Supplementary figures and images for "Molecular and functional diversity of distinct subpopulations of extracellular vesicles from stressed pancreatic beta cells: implications for autoimmunity"

### SupplFig1_Giri et al

#### Slide 1
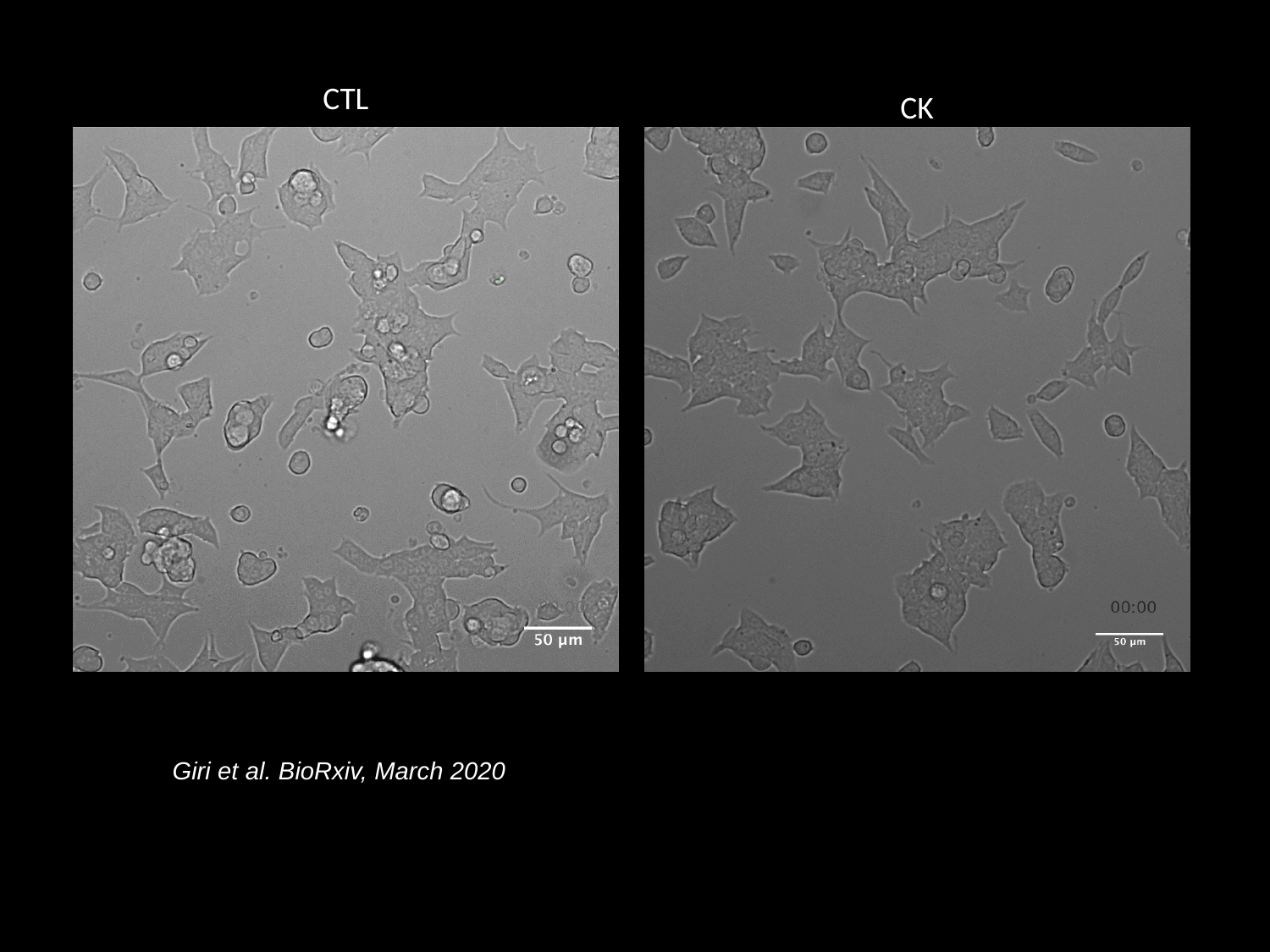

CTL
CK
Giri et al. BioRxiv, March 2020
